# Single cell evaluation of endocardial HAND2 gene regulatory networks reveals critical HAND2 dependent pathways impacting cardiac morphogenesis

**DOI:** 10.1101/2022.09.29.510167

**Authors:** Rajani M George, Beth A Firulli, Ram Podicheti, Douglas B Rusch, Brandon J Mannion, Len A Pennacchio, Marco Osterwalder, Anthony B Firulli

**Affiliations:** Herman B Wells Center for Pediatric Research, Departments of Pediatrics, Anatomy and Medical and Molecular Genetics, Indiana Medical School, Indianapolis, IN 46202; Center for Genomics and Bioinformatics, Indiana University, Bloomington, United States; Environmental Genomics and Systems Biology Division, Lawrence Berkeley National Laboratory, Berkeley, CA 94720, USA; Comparative Biochemistry Program, University of California, Berkeley, CA 94720, USA; US Department of Energy Joint Genome Institute, Lawrence Berkeley National Laboratory, Berkeley, CA 94720, USA; Department for BioMedical Research (DBMR), University of Bern, Bern, Switzerland; Department of Cardiology, Bern University Hospital, Bern, Switzerland

## Abstract

The transcription factor HAND2 plays critical roles during cardiogenesis. *Hand2* endocardial deletion (*H2CKO*) results in tricuspid atresia or double inlet left ventricle with accompanying intraventricular septum defects, hypo-trabeculated ventricles, and an increased density of coronary lumens. To understand the regulatory mechanisms of these phenotypes, single cell transcriptome analysis of E11.5 *H2CKO* hearts was performed revealing a number of disrupted endocardial regulatory pathways. Utilizing HAND2 DNA occupancy data, we identify several HAND2-dependent enhancers, including two endothelial enhancers for the sheer-stress master regulator, KLF2. A 1.8kb enhancer located 50kb upstream of the *Klf2* transcriptional start site imparts specific endothelial/endocardial expression within the vasculature and endocardium. This enhancer is HAND2-dependent for ventricular endocardium expression but HAND2-independent for *Klf2* vascular and valve expression. Deletion of this *Klf2* enhancer reveals reduced *Klf2* expression within ventricular endocardium. These data reveal that HAND2 functions within endocardial gene regulatory networks including sheer stress response.

## INTRODUCTION

Cardiac morphogenesis is a complex process requiring the synergistic action of multiple tissue types and fine-tuned control of morphogenetic events that are coordinated though the actions of transcription factors within each contributing cell type. Spatial and temporal specific cell signaling between the developing myocardium, the muscular portion of the heart, and endocardium, the inner endothelial lining of the heart, is required for normal cardiogenesis. The embryonic day (E)12.5 myocardium of the developing ventricles, expresses vascular endothelial growth factor A (VEGFA) which signals to the endocardium through its interactions with VEGF Receptor 2 (VEGFR2) establishing the coronary plexus, which will mature to contribute to the coronary arteries (Wu et al., 2012). In the atria, NOTCH signaling initiated within the endocardium communicates with receptors expressed in the myocardium regulating valve and sinoatrial node development (Y. Wang, Lu, Jiang, Wu, & Zhou, 2020; Y. Wang et al., 2013). Ventricular NOTCH signaling from the endocardium is also required for myocardial BMP10 expression which is essential for normal trabeculation (H. Chen et al., 2004; Del Monte-Nieto et al., 2018; Grego-Bessa et al., 2007).

Recent work has revealed that the basic helix loop helix (bHLH) transcription factor, HAND2 is required for Notch-dependent functions within the endocardium, modulating trabeculation, septation, coronary vascular maturation, as well as endocardial maturation within the embryonic heart (VanDusen, Casanovas, et al., 2014). Conditional endocardial deletion of *Hand2* using *Nfatc1*^*Cre*^ (*H2CKO*; (VanDusen, Casanovas, et al., 2014; Wu et al., 2012) results in embryonic lethality by E12.5. Embryos exhibit TA or DILV where both tricuspid and mitral valves connect the atria with the left ventricle (LV; (VanDusen, Casanovas, et al., 2014). *H2CKO*s also exhibit hypoplastic myocardium, an intraventricular septum (IVS) that is shifted to the right generating a smaller right ventricle (RV) and larger LV. *H2CKO* hearts are hypotrabeculated, and occasionally present with multiple IVS (VanDusen, Casanovas, et al., 2014). In addition to these defects, *H2CKO* hearts exhibit a pronounced hypervascularization phenotype with an increased number of coronary arteries within the myocardium (VanDusen, Casanovas, et al., 2014).

To investigate transcriptomic changes within *H2CKO*s endocardium, we employ single cell (sc) RNA seq to identify several gene regulatory networks (GRNs) compromised by HAND2 loss-of-function. GRNs impacted by loss of HAND2 include Wound Healing Signaling Pathway, Pulmonary Fibrosis Idiopathic Signaling, Pulmonary Healing Signaling, Tumor Microenvironment, NF-kb Signaling, STAT3 Pathway, and interestingly, the Apelin Endothelial Signaling Pathway related to shear stress response. Based on the overlap of significant gene expression changes with established HAND2 DNA occupancy data (Laurent et al., 2017), we sought to identify putative HAND2-dependent endocardial transcriptional enhancers. We selected 5 genes that showed robust expression changes associated with bound HAND2 to further evaluate: *Igf2, Igf2R, Ptn, Tmem108*, and *Klf2*. F0 transgenic reporter analysis reveals direct endocardial/endothelial transcriptional regulation of two of the selected genes, *Igf2R* and the shear stress master regulator gene *Klf2*. We further characterized the activity of a 1.7kb *Klf2* HAND2-binding conserved non-coding sequence located -23kb upstream of the *Klf2* transcriptional start site and another 1.8kb *Klf2* enhancer located -50kb upstream of the *Klf2* transcriptional start site (TSS). We observe that the -50kb *Klf2* enhancer is active within the early developing vasculature endothelium, and importantly, within the endocardium, recapitulating the endogenous *Klf2* expression pattern. To determine if HAND2 is necessary for *Klf2* enhancer endocardial expression *in vivo*, we interrogated the activity of the -50kb *Klf2* enhancer on the *H2CKO* background. Indeed, data reveal that in the absence of HAND2, activity of the -50kb *Klf2* enhancer is robustly reduced within trabecular endocardium but is unaffected within the systemic vasculature and developing valves. Gene edited deletion of the -50kb *Klf2* (*Klf2*^*Δ-50:(3*.*9kb)/Δ-50:(3*.*9kb)*^) enhancer shows that it is not required for viability. Gene expression analysis shows *Klf2* endocardial expression is significantly reduced by 40% suggesting that the -23kb *Klf2* enhancer or another yet to be identified enhancer are sufficient to maintain some *Klf2* endocardial expression. Collectively, these findings demonstrate several novel endocardial HAND2-dependent gene regulatory pathways including the shear stress response pathway, mediated in part, through HAND2 regulation of *Klf2*.

## RESULTS

### Deletion of Hand2 results in disruption of a number of endocardial GRNs including sheer stress pathway

To further investigate the role of HAND2 within the developing endocardium, we crossed *Hand2* conditional mice (*H2*^*fx/fx*^;*R26R*^*mTmG/mTmG*^, (Morikawa, D’Autreaux, Gershon, & Cserjesi, 2007; Muzumdar, Tasic, Miyamichi, Li, & Luo, 2007) with the endocardial specific *Nfatc1*^*Cre*^(Wu et al., 2012) to generate *H2CKO’s* (*Nfatc1*^*Cre*^*Hand2*^*fx/fx*^*R26R*^*mTmG/wt*^) and *Control* littermates (*Hand2*^*fx/+*^*R26R*^*mTmG/wt*^) for isolation of E11.5 hearts for single cell RNA-seq analysis on the 10x Genomics platform. In *H2CKO* hearts, *Cre-recombinase* mediated recombination leads to switching of tdtomato epifluorescence to that of *GFP* allowing for identification of embryos carrying the *Cre* allele. *Hand2* conditional allele status within the harvested litter is identified using rapid genotyping PCR. 13,885 unique barcodes were sequenced from a single *H2CKO* heart, and 14,259 barcodes were sequenced from a single *Control* heart. Based on high expression of hemoglobin genes, we excluded 5,828 barcodes from *H2CKO*s and 3,150 barcodes from *Control* hearts. Next, barcodes with total number of genes greater than 2500, indicating multiplets, were excluded from the analysis. We then utilized the remaining 5408 barcodes from *Control* and 6232 barcodes from *H2CKO*s for further analysis. Non-linear dimensionality reduction using uniform manifold approximation and projection (UMAP) plots resulted in 13 transcriptionally distinct clusters (Fig. 1A; Supplemental Spreadsheet 1).

**Figure 1:**
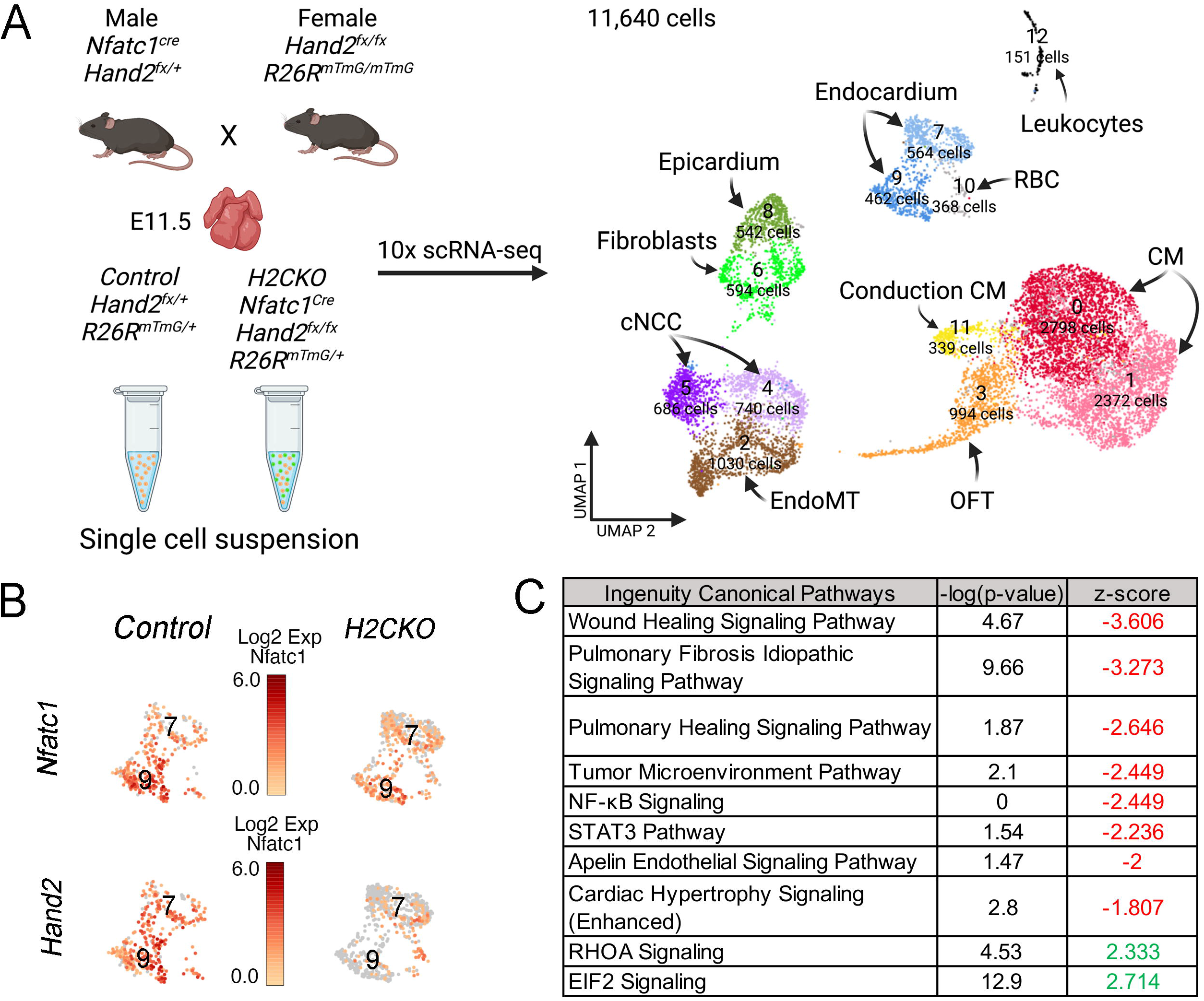
scRNA-seq analysis of *Nfatc1*^*cre*^ *Hand2*^*fx/fx*^ E11.5 ventricles. **A**. UMAP plot of all barcodes captured with single cell RNA-sequencing of E11.5 embryos from *Control* (*Hand2*^*fx/+*^R262R^mTmG/+^) and *H2CKO* (*Nfatc1*^*cre*^*Hand2*^*fx/fx*^R262R^mTmG/+^) hearts. *Control* n = 5408 *H2CKO* n = 6232 cNCC cardiac neural crest cells, CM cardiomyocytes, RBC red blood cells, OFT outflow tract mesenchyme, EndoMT endothelial to mesenchymal transition. **B-C**. Expression of *Hand2* and *Nfatc1* in endocardial clusters 7 and 9 in *Control* and *H2CKO*. **D**. IPA analysis of top 10% of differentially expressed genes in *Control* vs *H2CKO* populations within cluster 7. Z-scores in red represent downregulated pathways, z scores in green represent upregulated pathways.

Cluster identity is assigned by comparing gene expression of a gene in the *Control* cluster against expression of the same gene in all other *Control* clusters combined at a threshold of 0.25 log2FC, which establishes a rigorous threshold for significance (Supplemental Spreadsheet 1). Clusters 0 (red shaded) and 1 (pink shaded), representing 2793 cells, exhibit similar transcriptional profiles, and express cardiomyocyte marker gene transcripts, *Myh6* (99.4% cells in cluster 0, 98.1% cells in cluster 1) coding alpha myosin heavy chain (*α*MHC), and *Actinin alpha 2* (*Actn2*; 99.9% of cells in cluster 0 and 1; Fig. 1A; Supplemental Fig 1; Supplemental Spreadsheet 1). The 1030 cells within cluster 2 (brown shaded) express the extracellular matrix protein coding gene *Periostin* (*Postn*, 99.8% of cells), the bHLH transcription factor gene *Twist1* (99.1% of cells), the matricellular protein coding gene *transforming growth factor beta-induced* (*TGFBI*, 98% of cells) likely representing AV cushion cells that have undergone endothelial to mesenchymal transition (EMT, Supplemental Fig 1; Supplemental Spreadsheet 1). Cluster 3 (orange shaded) consists of 993 cells that we identify as outflow tract mesenchyme based on the expression of the neurovascular guiding factor *Semaphorin 3c* (87% of cells) and *Bone Morphogenetic Protein, Bmp4* (88% of cells; Supplemental Fig 1; Supplemental Spreadsheet 1). Cluster 4 light purple; 740 cells) and 5 (dark purple; 686 cells) consist of cardiac neural crest cells as determined by the expression of *Twist1* (99.8% of cells in cluster 4, 100% in cluster 5), *Insulin growth factor Igf1* (98% of cells in both clusters), and high mobility group transcription factor *Sox9* (94% of cells in cluster 4 and 96% of cells in cluster 5, Supplemental Fig 1; Supplemental Spreadsheet 1). The 594 cells in cluster 6 are epicardial cells (light green shaded), undergoing transition to a fibroblast phenotype as marked by expression of the growth factor *Pleiotrophin* (*Ptn*,100% of cells, T-box transcription factor *Tbx18* (90% of cells), and bHLH transcription factor *Tcf21* (98% of cells; Supplemental Fig 1; Supplemental Spreadsheet 1). Cluster 7 contains 564 cells (light blue shaded) and are defined as endocardial by the expression of the transmembrane transport protein *Ramp2* (100% of cells), the *vascular endothelial cadherin Cdh5* (99% of cells, Supplemental Fig 1; Supplemental Spreadsheet 1), *Platelet endothelial cell adhesion molecule Pecam1* (95% of cells), and the nuclear factor of activated T cells *Nfatc1* (64% of cells). Cluster 8 consists of 542 epicardial cells (dark green shaded) expressing *Ptn* (100% of cells), *Tbx18* (97% of cells), and *Tcf21* (97% of cells; Supplemental Fig 1; Supplemental Spreadsheet 1). Cluster 9 represents a second endocardial cell cluster (dark blue shaded) wherein 462 cells express *Ramp2* (100% of cells), *Cdh5* (95% of cells), *Pecam1* (98% of cells), and *Nfatc1* (88% of cells; Supplemental Fig 1; Supplemental Spreadsheet 1). Cluster 10 represents blood cells populations (grey shaded) marked by hemoglobin (*Hbb*) expression in 100% of cells. Cluster 11 appears to represent conduction system (yellow shaded) marked by expression of Calcium channel, voltage-dependent, *α*2/*δ*2 *subunit 2 Cacnα2δ2* (97% of cells) and *Calcium channel, voltage-dependent, T type, α 1H subunit, Cacnα1h* (65% of cells). Cluster 12 (black shaded) is composed of lymphocytes indicated by expression of *interferon induced transmembrane protein Ifitm3* (47% of cells) *and Histocompatibility 2, D region locus 1 H2-D1* (93% of cells; Supplemental Fig 1; Supplemental Spreadsheet 1). *Nfatc1* expression robustly marks both cluster 7 (64% of cells) and 9 (88% of cells; Fig. 1B). Comparison of *H2CKO* and *Control* barcodes specific for *Hand2* expression exhibit robust downregulation within these two endocardial clusters in the presence of *Nfatc1*^*Cre*^ (cluster 7 log2FC -1.5 *p* = 1.5×10^−52^, cluster 9 log2FC -1.66 *p* = 9.3×10^−55^, Fig. 1B, Supplemental Fig 2).

### Analysis of endocardial clusters 7 and 9 indicates misregulation within several endocardial gene regulatory networks

To examine transcriptome data in an unbiased fashion, we employed Ingenuity Pathway Analysis (IPA) on differentially expressed genes (log2FC>±0.5) from *H2CKO*s and *Control* cells (Fig. 1C, Supplemental Fig 3, Supplemental Spreadsheet 3). Loss of *Hand2* within the endocardium leads to significant changes in developmental, morphological, and cardiovascular gene regulatory networks (Fig 1C, Supplemental Fig 4). IPA on cluster 7 indicates that *Hand2* is downregulated within the Cardiac Hypertrophy Signaling (Enhanced) canonical pathway (z-score -1.807, -log p-value 2.8, Supplemental Spreadsheet 3), which is close to a significant z-score absolute value of 2.

We also observe abnormal expression of a number of endocardial transcripts: *Endothelin converting enzyme* 1 (*Ece1*), a sheer stress responsive gene expressed by vascular endothelial cells and required for formation of patent vasculature in the developing heart (Masatsugu et al., 2003; Robinson et al., 2014). *Ece1* is significantly downregulated within *H2CKO* endocardium (cluster 7 log2FC -1.16 *p* = 2.69×10^−28^; cluster 9 log2FC -0.45 *p* = 9.05×10^4^; Supplemental Spreadsheet 2). Concomitantly, *Ece1*’s substrate, *Endothelin1* (*Edn1*), a potent vasoconstrictor that is secreted by endothelial cells when laminar flow induces sheer stress (Morawietz et al., 2000), is upregulated within *H2CKO* endocardium (cluster 7 log2FC 0.58 *p* = 1.9×10^−9^). Previous work shows that EDN1 signaling lies upstream of *Hand2* within the cranial neural crest cells during craniofacial morphogenesis (Charite et al., 2001; Clouthier et al., 2000). Thus, the observed increase in *Edn1* could reflect feedback compensation resulting from of endocardial loss of *Hand2*.

*Fibronectin* (*Fn1*), a component of the extracellular matrix secreted by endothelial cells, is significantly downregulated within clusters 7 and 9 cell analysis (cluster 7 log2FC -1.17, *p* = 1.4×10^−21^; cluster 9 log2FC -0.66, *p* = 1.6×10^−4^, Supplemental Spreadsheet 2). *Fn1* is listed in multiple pathways in IPA (Fig. 1C) including: Wound Healing Signaling Pathway (z-score -3.606, -log p-value 4.67), Pulmonary Fibrosis Idiopathic Signaling Pathway (z-score -3.273, -log p-value 9.66), and Tumor Microenvironment Pathway (z-score -2.449, -log p-value 2.1).

The mechanosensitive transcription factor *hypoxia inducible factor 1 alpha* (*Hif1α*) is significantly down regulated within endocardial clusters (cluster 7 log2FC -0.58 *p* = 4.5×10^−12^; cluster 9 log2FC -0.48 *p* = 2.3×10^−10^, Supplemental Spreadsheet 2, (Feng, Fragiadaki, Souilhol, Ridger, & Evans, 2017). IPA reveals related canonical pathways that include *Hif1*α are disrupted (Fig. 1C, Supplemental Spreadsheet 5): Pulmonary Healing Signaling Pathway (z-score -2.65, - log p-value 1.87), and Tumor Microenvironment Pathway (z-score of -2.449, -log *p* value 2.1). The Tumor Microenvironment pathways also include the gene that codes for the insulin growth factor, *Igf2*, which is also significantly down regulated in *H2CKO* endocardial clusters (cluster 7 log2FC -1.44 *p* = 3.2×10^−46^, cluster 9 log2FC -1.81 *p* = 2.1×10^−65^). Indeed, *HIF-1*α transcriptionally regulates *Igf2* via hypoxia responsive elements at the *Igf2* locus (Feldser et al., 1999). Within endocardial cells, angiogenesis requires the action of the sheer stress master regulator KLF2 (Nigro, Abe, & Berk, 2011). Coronary angiogenesis initiates at E11.5 with formation of mature coronary plexus by E13.5 (Tian et al., 2013). Lineage tracing experiments have shown that a major source of the coronaries are the sinus venosus and endocardium with some contribution from the epicardium (Rhee et al., 2021). KLF2 expression within endocardial cells exerts an anti-angiogenic effect by modulation of *VegfR2/Kdr* expression by KLF2 binding at the promoter region (Bhattacharya et al., 2005). KLF4 is expressed within these same populations of cells (Jain, Sangwung, & Hamik, 2014); however, it’s precise role in sheer stress response remains unclear (Sangwung et al., 2017).

### The shear stress master regulator gene Klf2 is specifically downregulated within ventricular endocardium in H2CKO endocardium

IPA also reveals that *H2CKO*s exhibit a significant downregulation of the Apelin Signaling Pathway with z score of -2 (−log p value 1.47, Fig. 1C). Apelin is an angiogenic factor that controls migration of endothelial cells and is required for the normal development of blood vessels (Helker et al., 2020; Kwon et al., 2016; L. Lu, Wu, Li, & Chen, 2017). The Apelin Signaling IPA pathway includes a major contributing factor to normal vasculogenesis and ventricular morphogenesis within the embryonic heart - sheer stress signaling (Haack & Abdelilah-Seyfried, 2016). We observe that the gene coding the sheer stress regulated transmembrane receptor *Heart of Glass* (*Heg1*) is significantly downregulated within both endocardial clusters (cluster 7 log2FC -0.36 *p* = 6.35×10^−9^, cluster 9 log2FC -0.9 *p* = 1×10^−15^, Supplemental Spreadsheet 2). *Heg1* zebrafish mutants exhibit significant vascular malformations (Kleaveland et al., 2009). Interestingly, the transcription factor KLF2 is a direct regulator of *Heg1* expression (Razani et al., 2001; Zhou et al., 2016). KLF2 is a well-studied sheer stress response transcription factor considered the master regulator of this response (Bhattacharya et al., 2005; Chiplunkar et al., 2013; Sangwung et al., 2017).

Since the hypervascularization phenotype observed in the *H2CKO*s could be caused by defective angiogenesis and KLF2 is a major regulator of angiogenesis, we employed *in-situ* hybridization to closely examine both *Hand2* and *Klf2* transcripts within E12.5 *Control* and *H2CKO* embryo hearts (Fig. 2). Results show that *Hand2* expression within the trabecular endocardium of the ventricles is significantly reduced in *H2CKOs* when compared to *controls* (Fig. 2A and C). *Klf2* is also robustly expressed within the ventricular endocardium (Fig. 2B’ black arrowheads), and particularly within areas of high sheer-stress such as the endocardial lining of the AV canal (Fig. 2B’ red arrowheads; (Chiplunkar et al., 2013). We observe that *Klf2* expression within the ventricular endocardium of *H2CKO*s is greatly reduced; however, *Klf2* expression within the endocardial lining of the AV canal as well as within the systemic vasculature is maintained (Fig. 2D’ compare black and red arrows). Next, we examined genes that are downstream of KLF2 that are significantly changed with *H2CKO* endocardial clusters 7 and 9 (Supplemental Table 1). Indeed, we find significant changes in gene expression within KLF2 target genes suggesting that loss of HAND2 in the endocardium reduces *Klf2* expression as well the expression of its targets (Supplemental Table 1).

**Figure 2:**
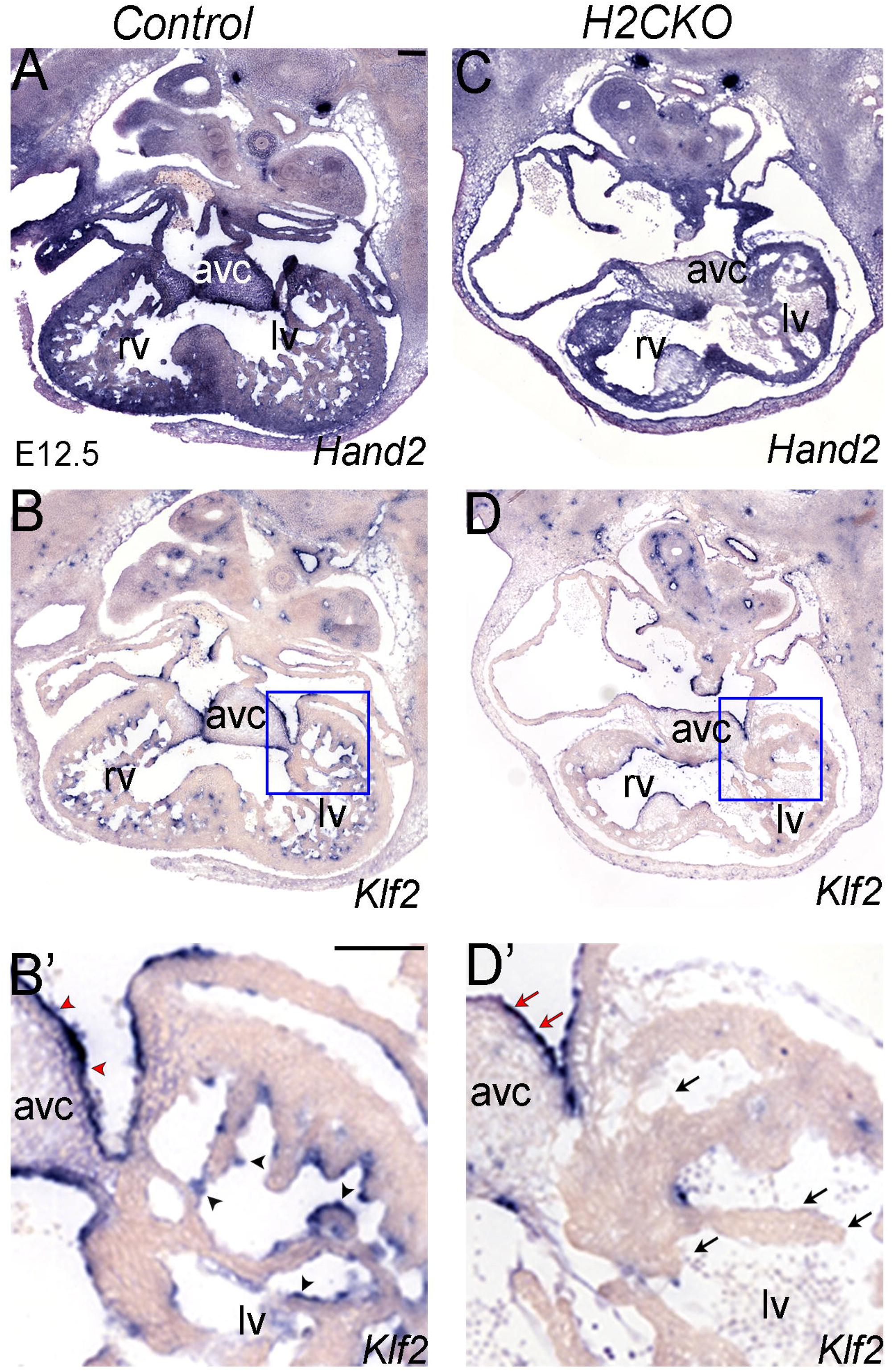
Expression of *Klf2* in *H2CKO*s. **A, B, B’**. Section *in-situ* hybridization showing *Hand2* and *Klf2* expression in E12.5 *Hand2*^*fx/fx*^ *controls*. Black arrowheads indicate ventricular endocardium. Red arrowheads indicate endocardium covering the AV cushion. n = 10. rv, right ventricle; lv, left ventricle, avc, atrioventricular canal. Scale bars 100µm. **C, D, D’**. Section *in-situ* hybridization showing *Hand2* and *Klf2* expression in E12.5 *Nfatc1*^*cre*^ *Hand2*^*fx/fx*^ *H2CKOs*. Black arrows indicate ventricular endocardium. Red arrows indicate endocardium covering the AV cushion.

**Table 1:**
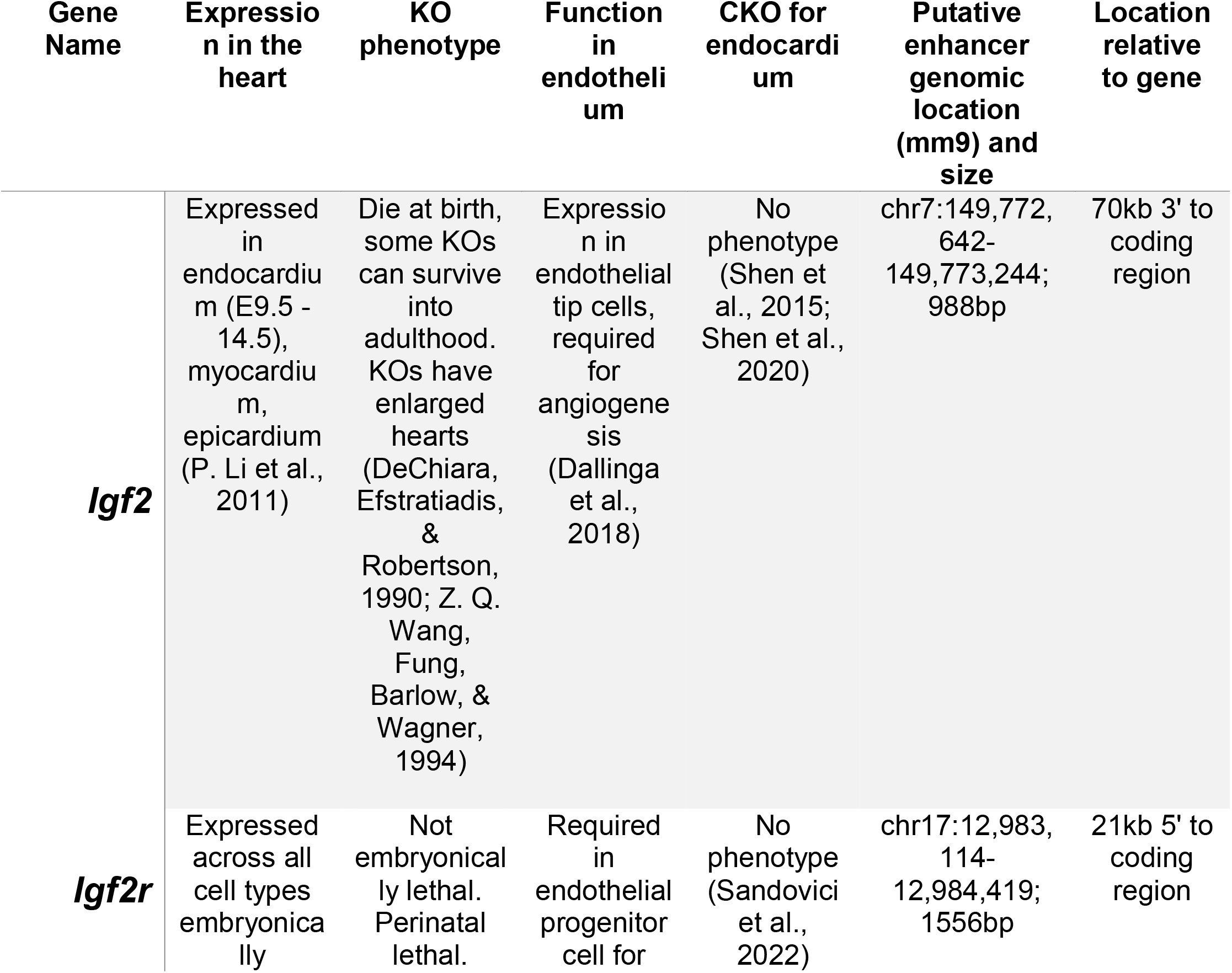
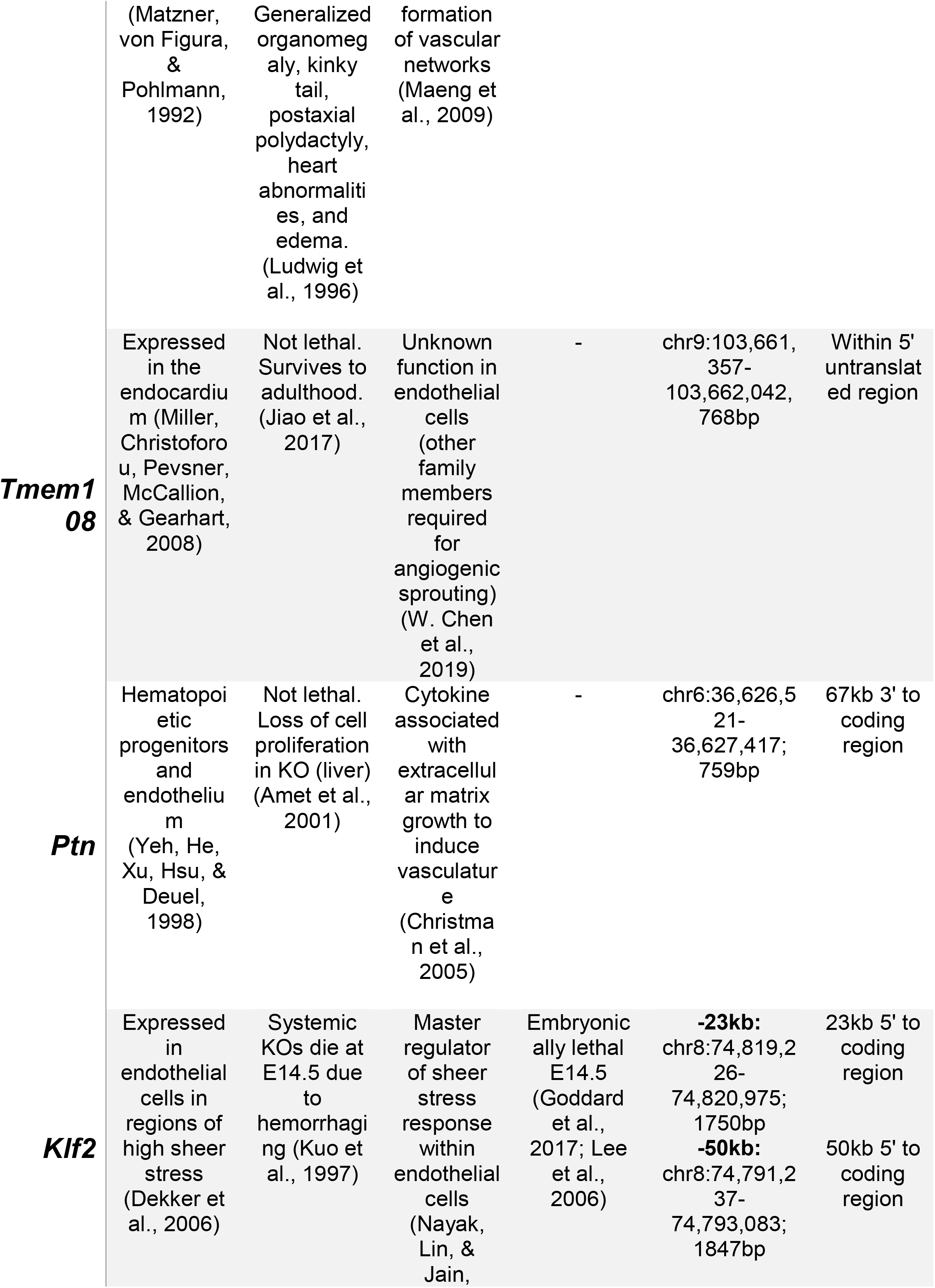
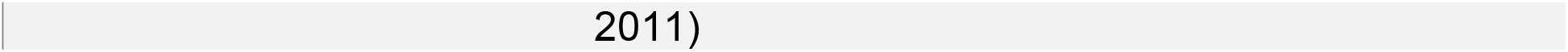
Regulated target genes that exhibit HAND2 DNA occupancy within CNS’s evaluated for enhancer activity.

### Comparison of scRNA-seq regulation and HAND2 DNA occupancy identifies three novel endothelial/endocardial enhancers

The differential gene expression profiles in endocardial clusters from *WT* and *H2CKO* hearts suggest direct interaction of the HAND2 transcription factor with *cis*-regulatory elements in the respective loci for transcriptional control. We selected a set of genes (*Igf2, Tmem108, Ptn, Igf2R*, and *Klf2*) and indeed distinct HAND2 interaction peaks were identified in the respective regulatory domains by determining regions of evolutionary conservation and HAND2 DNA binding (Fig. 4, Table 1). For identification of HAND2 target regions, we utilized established E10.5 chromatin immunoprecipitation ChIP-seq data from mouse embryonic hearts expressing a *Hand2*^*3xFlag*^ knock-in allele (Laurent et al., 2017). To validate and define the *in vivo* enhancer identity of these six putative enhancer elements we employed enSERT, a method for CRISPR-mediated site-directed reporter transgenesis targeting the *H11* locus (Kvon et al., 2020). Following PCR amplification from murine genomic DNA, we cloned the selected elements into an enSERT vector containing a human *beta-globin* promoter-driven *β-galactosidase* (LacZ) cassette.

Three of our putative HAND2-dependent enhancers that exhibit evolutionary conservation and HAND2 DNA occupancy did not exhibit endocardial/endothelial enhancer activity (Supplemental Fig 5). IGF2 is a secreted growth factor expressed within the epicardium and endocardium during heart development (Shen et al., 2015) and is highly down regulated within clusters 7 and 9 in *H2CKO*s (Supplemental Spreadsheet 2). At the *Igf2* genomic locus, a 988 bp CNS located 70kb 3’ to the coding region (Supplemental Fig 5A) contained a conserved non-coding element with HAND2 DNA occupancy. However, analysis by enSERT exhibited no enhancer activity of this element (n=9; Supplemental Fig 5B and B’). TMEM108 is implicated as a marker for progenitor epicardial cell populations although its role within the endocardium is currently unclear (Bochmann et al., 2010). *Tmem108* is significantly down regulated within endocardial clusters 7 and 9 (Supplemental Spreadsheet 2); however, a HAND2-occupied 768bp CNS 5’ of *Tmem108* exhibited no enhancer activity (n=3; Supplemental Fig 5C, D and D’). *Pleiotrophin* (*Ptn*) codes for a secreted cytokine, is an inducer of EMT, and is mitogenic to endothelial cells, resulting in angiogenesis (Perez-Pinera, Berenson, & Deuel, 2008). *Ptn* is significantly down regulated within both cluster 7 and 9 (Supplemental Spreadsheet 2). We tested a HAND2-occupied 759bp CNS 3’ of *Ptn*; however, results show no E11.5 heart expression (n=3; Supplemental Fig 5E, F and F’)

Three of our putative HAND2-dependent enhancers that exhibit robust HAND2 DNA occupancy did exhibit endocardial/endothelial enhancer activity. We successfully interrogated a 1556 bp IGF receptor 2R (*Igf2R*) HAND2-occupied CNS conserved element 21kb 5’ of its transcriptional start site (Fig. 3A). *Igf2R* is robustly expressed within the endocardium (K. Wang et al., 2019) and lacks a tyrosine kinase domain acting as a negative regulator of IGF2 (Braulke, 1999; Ludwig et al., 1996). *Igf2r* is significantly downregulated within *H2CKO*s endocardial clusters 7 and 9 (Supplemental Spreadsheet 2). F0 analysis of the *Igf2R* HAND2-occupied CNS generated seven E11.5 embryos out of eight that exhibit endocardial specific *lacZ* staining representing a novel endocardial enhancer (Fig. 3B and B’).

**Figure 3:**
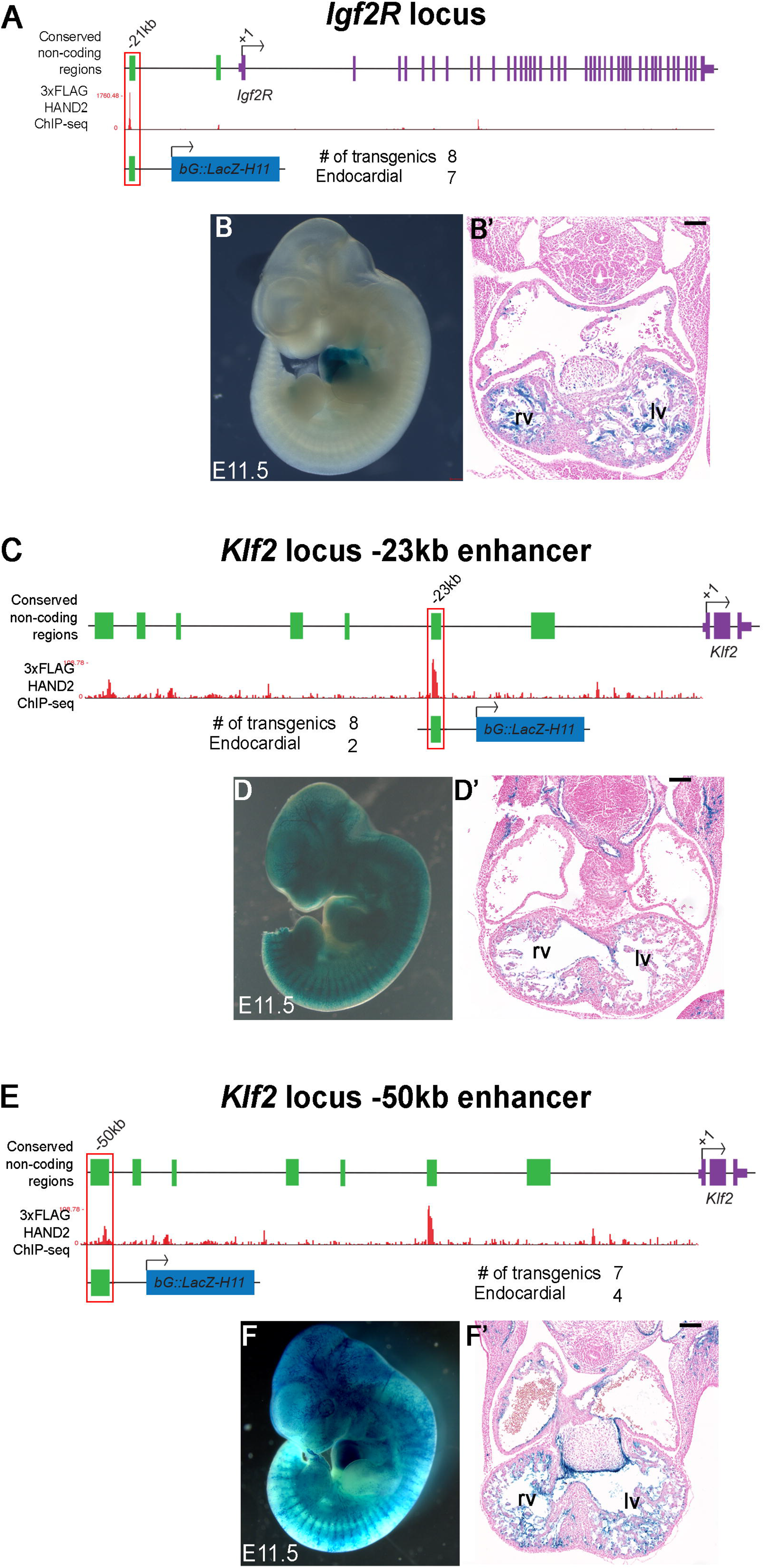
F0 reporter expression analysis of target genes showing both altered gene expression and HAND2 DNA-occupancy. **A**. *Igf2R* genomic locus showing conserved non-coding regions (green solid boxes), transcriptional start site (+1), relative location of enhancer element (−21kb, red outline). HAND2-3xFlag ChIP-seq data (Laurent et al., 2017) showing genomic regions of HAND2 binding. **B, B’**. -21kb HAND2 binding conserved non-coding region at *Igf2R* locus used to make transgenic F0 embryos and *lacZ* staining results. Representative whole mount image of E11.5 transgenic embryo. Numbers of transgenic F0 embryos obtained, 2. Numbers of F0 embryos that showed endocardial staining, 2. **C**. *Klf2* genomic locus showing conserved non-coding regions (green solid boxes), transcriptional start site (+1), relative location of enhancer element (−23kb, red outline). HAND2^3xFlag^ ChIP-seq data (Laurent et al., 2017) showing genomic regions of HAND2 binding. **D. D’**. -23kb HAND2 binding conserved non-coding region at *Klf2* locus used to make transgenic F0 embryos and *lacZ* staining results. Representative whole mount image of E11.5 transgenic embryo. Numbers of transgenic F0 embryos obtained, 8. Numbers of F0 embryos that showed endothelial/endocardial staining, 2. **E**. *Klf2* genomic locus showing conserved non-coding regions (green solid boxes), transcriptional start site (+1), relative location of enhancer element (−50kb, red outline). HAND2^3xFlag^ ChIP-seq data (Laurent et al., 2017) showing genomic regions of HAND2 binding. **F. F’**. -50kb HAND2 binding conserved non-coding region at *Klf2* locus used to make transgenic F0 embryos and *lacZ* staining results. Representative whole mount image of E11.5 transgenic embryo. Numbers of transgenic F0 embryos obtained, 7. Numbers of F0 embryos that showed endothelial/endocardial staining, 4. Scale bar 100µm. lv left ventricle, rv right ventricle.

From the observation that both *Igf2R* and *Igf2* are both downregulated within the endocardial clusters, we wanted to determine if endocardial proliferation was affected in *H2CKOs*. We conducted differential abundance analysis to determine cell type representation within *H2CKO*s and *control* barcodes (Supplemental Table. 2). Results indicate a significant increase in the number of barcodes in endocardial population (cluster 7), as well as increase in cardiomyocyte population (clusters 0 and 1), suggesting that loss of IGF2R might affect cell numbers in *H2CKO*s. The discovery of a HAND2-binding CNS -21kb 5’ of the *Igf2r* TSS that drives endocardial-specific reporter expression supports this idea (Fig. 3A, B and B’). In order to determine the evolutionary conservation of this element, we utilized CLUSTWAL analysis, which indicates conservation within rodents but not broadly within mammals (Supplemental Fig. 6).

### Two conserved non-coding elements drive expression of Klf2 in ventricular endocardium

We next interrogated two HAND2-occupied CNS within the shear stress master regulator KLF2. Expression analysis in both the scRNA-seq analysis and *in situ* hybridization experiments reveal dynamic *Klf2* regulation within the heart (Fig. 2B, B’, D and D’). *Klf2* expression is specifically downregulated regulated by HAND2 within the ventricular endocardium; however, *Klf2* endothelial expression is maintained within AV cushion endocardium (Fig. 2B’, D’). This observation is consistent with the downregulation of *Hand2* expression within the AV cushion endocardium post EMT as previously reported (VanDusen, Casanovas, et al., 2014) and suggests HAND2-independent regulation of *Klf2* transcription within the ventricular and AV cushion endocardium, as well as the systemic vasculature where *Hand2* is not expressed (VanDusen, Casanovas, et al., 2014).

Previously, a CNS located 100bp upstream of the *Klf2* transcriptional start site has been shown to be responsive to sheer stress (Huddleson, Srinivasan, Ahmad, & Lingrel, 2004), but HAND2 DNA occupancy data does not indicate HAND2 DNA-binding within this element (Laurent et al., 2017). Interestingly, we identified two HAND2-occupied CNS within *Klf2*, one (1.7kb) located -23kb upstream of the *Klf2* TSS and another CNS (1.8kb) located -50kb relative to the *Klf2* TSS (Fig. 3C-F). The -23kb *Klf2* CNS exhibits high level of HAND2 occupancy and sequence conservation (Fig. 3C). Results show 2 out of 8 F0 transgenics exhibit *lacZ* staining within embryonic vasculature and endocardium (Fig. 3D and D’). Although encouraging, six transgenic littermates did not show staining in a consistent pattern; 4 embryos revealed no visible staining, 3 embryos have staining only within the vasculature, 3 embryos exhibit staining within the AV canal, where two of these exhibited some endocardial staining within ventricles (data not shown). Out of these two, we observe only one embryo showing consistent staining throughout the left and right ventricular endocardium. The -50kb *Klf2* CNS also exhibits robust HAND2 DNA occupancy and sequence conservation (Fig. 3E) and was used to generate E11.5 F0 transgenics. Out of 7 F0 transgenics generated, 4 exhibited robust *lacZ* staining within the endocardium and systemic vasculature (Fig. 3F and F’).

Given the higher consistency in exhibiting endocardial/endothelial activity from the -50kb *Klf2* enhancer element (57% of F0s), we generated stable *lacZ* transgenic lines using the -50kb CNS (Fig. 4A). At E11.5, out of 12 transgenic lines generated, 5 (41%) recapitulated consistent and robust *lacZ* staining within the endocardium and systemic vasculature (Supplemental Fig. 7). The remaining 7 lines exhibited no observable E11.5 *lacZ* staining. We next examined additional embryonic time points using a single line (Line # 901, Supplemental Fig. 7). Analysis of reporter activity in E7.5 embryos reveals *lacZ* staining within endothelial precursors, the blood islands (Fig. 4B), and within the dorsal aorta at E8.5 (Fig. 4C). At E9.5 and E10.5, the -50kb CNS robustly drives *β-galactosidase* expression within endothelial structures, within the branchial arches, and intersomitic blood vessels (Fig. 4D and E arrow). Histological cross sections of *lacZ* stained torsos counter stained with NFR at E10.5 reveals endocardial specific staining at this time point (Fig. 4F, F’, G and G’). Thus, the -50kb *Klf2* CNS acts as a transcriptional enhancer within endothelial cells of developing vasculature including the endocardium and AV cushions (Fig. 4F’ and G’).

**Figure 4:**
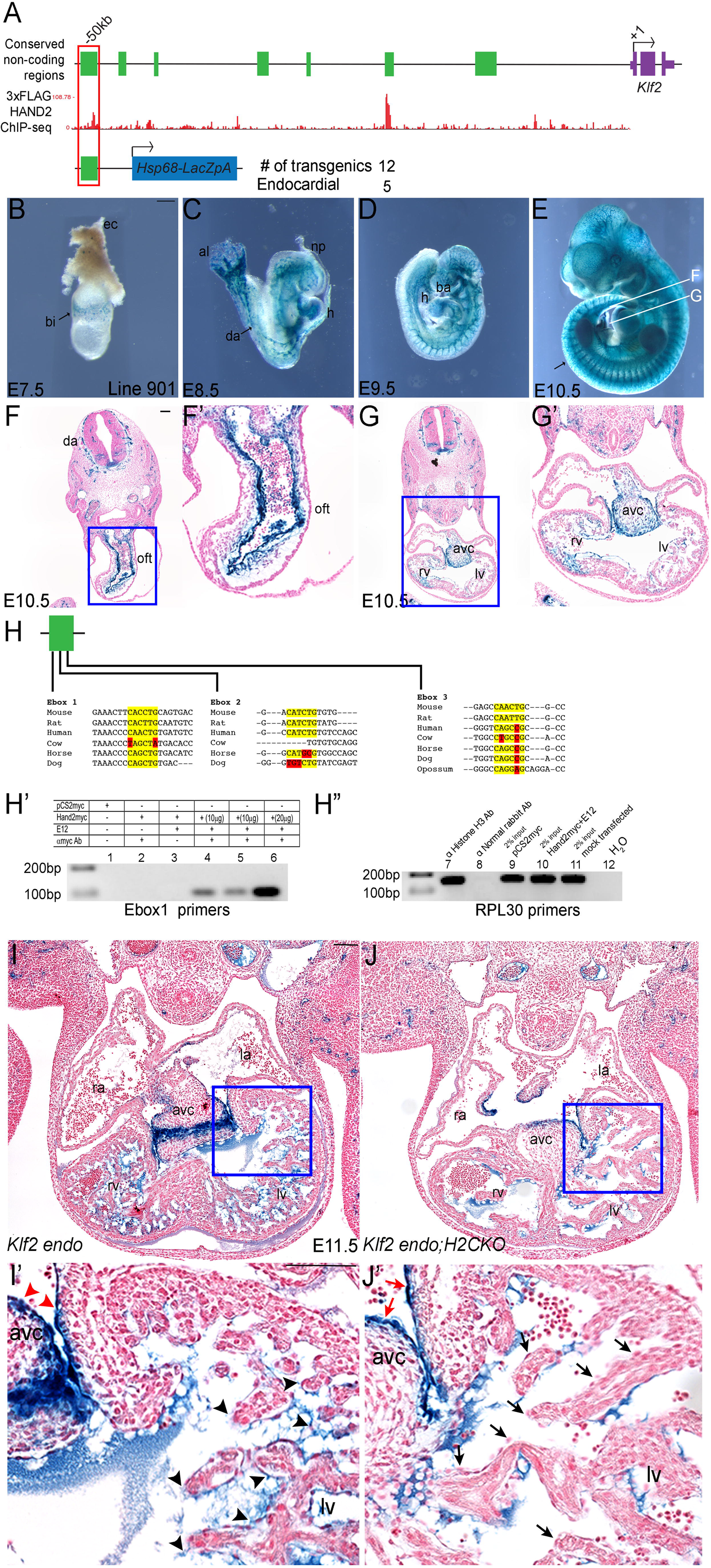
HAND2 responsive conserved *Klf2* enhancer is HAND2 dependent. **A**. Conserved non-coding regions (green boxes) upstream of *Klf2* transcriptional start site (+1). HAND2^3xFlag^ ChIP-seq data (Laurent et al., 2017) showing regions of *Hand2* binding. **B-E**. -50kb HAND2-binding conserved non-coding *Klf2* region used to make stable transgenics, numbers of stable transgenic lines obtained, and number of lines showing endocardial staining of transgene. *lacZ* staining of E7.5, E8.5, E9.5, E10.5 embryos from founder line 901. Plane of cross section for outflow tract (oft) and four chamber view are indicated by white lines in E. bi, blood islands; ec, ectoplacental cone; al, allantois; h, heart; np, neural plexus; da, dorsal aorta; ba, branchial arches. Arrow in panel E indicates inter-somitic blood vessels. Scale bar in B is 200µm. **F F’**. Transverse sections counterstained with Nuclear Fast Red (NFR) showing *lacZ* staining in the outflow tract. da, dorsal aorta; oft, outflow tract. Blue box in F indicates area of zoom in F’. Scale bar 100µm. **G G’**. Transverse sections counterstained with NFR showing *lacZ* staining in four chamber view. avc, atrioventicular cushion; rv, right ventricle; lv, left ventricle. Blue box in G indicates area of zoom in G’. **H**. Conserved E-boxes in -50kb *Klf2* enhancer. Yellow basepairs indicate regions of conservation. Red base pairs indicate regions of non-conservation within the canonical E-box sequence. **H’**. ChIP experiment in NIH 3T3s transfected with plasmids as indicated. Primers specific for E-box1 showed binding when *mycHand2* construct is co-transfected with *E12* in dose dependent manner. 1. pCS2myc empty vector; 2. pCSmyc; αmycAb; 3. mycHand2; No Ab; 4. 10µg mycHand2+E12; αmycAb; 5. 10µg mycHand2+E12; αmycAb; 6. 20µg Hand2+E12; αmycAb **H”**. Control primers against mouse RPL30 show used to show control ChIP using anti-HistoneH3 antibody and 2% input samples as indicated. Negative control using ChIP with anti-Normal rabbit antibody does not show signal. Primers used for PCR specific for mouse RPL30 gene intron 2. 7. αHistone H3 Ab; +ve cntl; 8. αNormal rabbit Ab; -ve cntl; 9. 2% input; pCS2myc; αHistone H3 Ab; 10. 2% input; mycHand2+E12; αHistone H3 Ab; 11. 2% input; mock transfected; αHistone H3 Ab; 12. H_2_O **I and I’**: -50kb *Klf2* enhancer at E11.5 were stained for X-gal, sectioned, and counter stained with NFR. Blue box in I indicates area of zoom in I’. Scale bar 100µm. Black arrowheads indicate ventricular endocardium with *lacZ* staining. Red arrowheads indicate *lacZ* staining within the endocardium covering the AV cushion. **J and J’**: *Nfatc1*^*Cre*^ *Hand2*^*fx/fx*^ with *Klf2* enhancer transgene at E11.5. Blue box in J indicates area of zoom in J’. Black arrows indicate loss of enhancer activity within the ventricular endocardium. Red arrows indicate *lacZ* staining within the endocardium covering the AV cushion. ra, right atria; la, left atria; rv, right ventricle; lv, left ventricle; avc, atrioventricular canal.

Motif analysis of the -50kb *Klf2* enhancer shows the presence of 3 conserved E-boxes (Fig. 4H, Supplemental Fig. 8) that lie within this established HAND2 DNA occupancy peak (Laurent et al., 2017). In order to determine the interaction between HAND2 and the 3-conserved E-boxes within this *Klf2* enhancer, we conducted ChIP assays in NIH-3T3 cells by co-transfecting plasmids encoding a 5’ Myc-tagged *Hand2* and an untagged *E12* (Fig. 4H’). Negative controls used pCS2+myc samples immunoprecipitated with and without *α*Myc, and Myc-*Hand2* immunoprecipitated without *α*Myc. ChIP-PCR, utilizing PCR primers specific for a 93bp region containing the 5’ most Ebox (Ebox1 CACCT), we observe HAND2 DNA binding at Ebox1 via with a dose dependent response in binding at the *Klf2* enhancer (Fig. 4H’). The controls for this experiment employ primers recognizing the mouse RPL30 gene (Fig. 4H”). Interrogation of HAND2 binding Ebox 2 and Ebox 3 revealed no DNA binding activity (not shown). These *in vitro* data support the DNA-occupancy and *Klf2* expression data that HAND2 directly binds and transcriptionally regulates *Klf2* through the -50kb *Klf2* CNS.

### HAND2 directly regulates expression of Klf2 within the ventricular endocardium

To assess if the endocardial -50kb *Klf2* enhancer is dependent on HAND2 *in vivo*, we crossed the -50kb *Klf2* enhancer *lacZ* reporter transgenic with the endocardial specific *H2CKO*. E11.5 embryos were *lacZ* stained, sectioned, and counter stained with NFR (Fig. 4I). -50kb *Klf2* enhancer embryos (*lacZ+ Hand2*^*fx/+*^) exhibit positive staining of trabecular endocardium (Fig. 4I and I’). In comparison, -50kb *Klf2* enhancer *lacZ* reporter *H2CKO* embryos (*lacZ+*; *Nfatc1*^*cre*^ *Hand2*^*fx/fx*^) show robust reduction in ventricular endocardial lacZ staining (Fig. 4J and J’, arrows) where *lacZ* staining within the endocardium over the developing AV cushions, and systemic vasculature is maintained (Fig. 4J).

The -50kb *Klf2* enhancer activity recapitulates the *Klf2* mRNA expression pattern throughout the embryo, with activity within the developing vasculature and the endocardium; however, crossing the -50kb *Klf2* enhancer *lacZ* reporter to the *H2CKO* background only alters enhancer *lacZ* staining within the ventricular endocardium and does not appreciably alter expression within the systemic vasculature or within the AV cushions, a result completely consistent with *in situ* hybridization analysis of *Klf2* expression in the *H2CKO* (Fig. 2B’, D’). These data suggest that HAND2 DNA binding within this -50kb *Klf2* endothelial/endocardial enhancer is necessary for its activity within the ventricular endocardium.

### *Deletion of the -50kb Klf2 endothelial/endocardial* enhancer results in decreased *Klf2* ventricular endocardial expression

As the -50kb *Klf2* endothelial/endocardial enhancer recapitulates *Klf2* vascular expression, we next tested its requirement for maintenance of *Klf2* expression via CRISPR-mediated genomic deletion in mice (Supplemental Fig. 9A). Eleven enhancer-deleted lines were obtained and 4 of these were crossed two generations with wildtype mice before they were intercrossed for homozygosity. *Klf2*^*Δ-50:(3*.*9kb)/Δ-50:(3*.*9kb)*^ mice were viable and were born at mendelian frequencies in four outcrossed founder lines (Supplemental Fig. 9B). A single line was then set up for timed pregnancies and E11.5 embryos were evaluated for *Klf2* expression (Fig. 5A and B). *Klf2* expression is visibly lower within the endocardium (Fig. 3 and 6). In contrast Hand2 expression within adjacent sections appears unchanged between controls and *Klf2*^*Δ-50:(3*.*9kb)/Δ-50:(3*.*9kb)*^ hearts (Fig. 5 C and D). To confirm that the observed *Klf2* expression drop in the *Klf2*^*Δ-50:(3*.*9kb)/Δ-50:(3*.*9kb)*^ homozygous hearts is significant, we dissected E11.5 ventricles from 8 *Klf2*^*Δ-50:(3*.*9kb)/Δ-50:(3*.*9kb)*^, and 10 wild type controls and performed qRTPCR for *Hand2* and *Klf2* (Fig. 5E). As predicted by the ISH analysis, *Klf2* expression levels are significantly lower (p< 0.001) in *Klf2*^*Δ-50:(3*.*9kb)/Δ-50:(3*.*9kb*^ ventricles when compared to controls ventricles, approximately 60% of what is observed in wild type (Fig. 5E). Expression results show that *Hand2* expression is largely unchanged within *Klf2*^*Δ-50:(3*.*9kb)/Δ-50:(3*.*9kb*^ hearts when compared to wild type controls (Fig. 5E). Given that the -50kb enhancer recapitulates all *Klf2* embryonic expression but is only affected by HAND2 within the ventricular endocardium, when we consider that two additional endothelial *Klf2* enhancers are unaltered (the -23kb *Klf2* CNS, and the proximal sheer stress responsive CNS previously reported (Huddleson et al., 2004), the observed 40% decrease in endocardial expression is in line with these observations.

**Figure 5:**
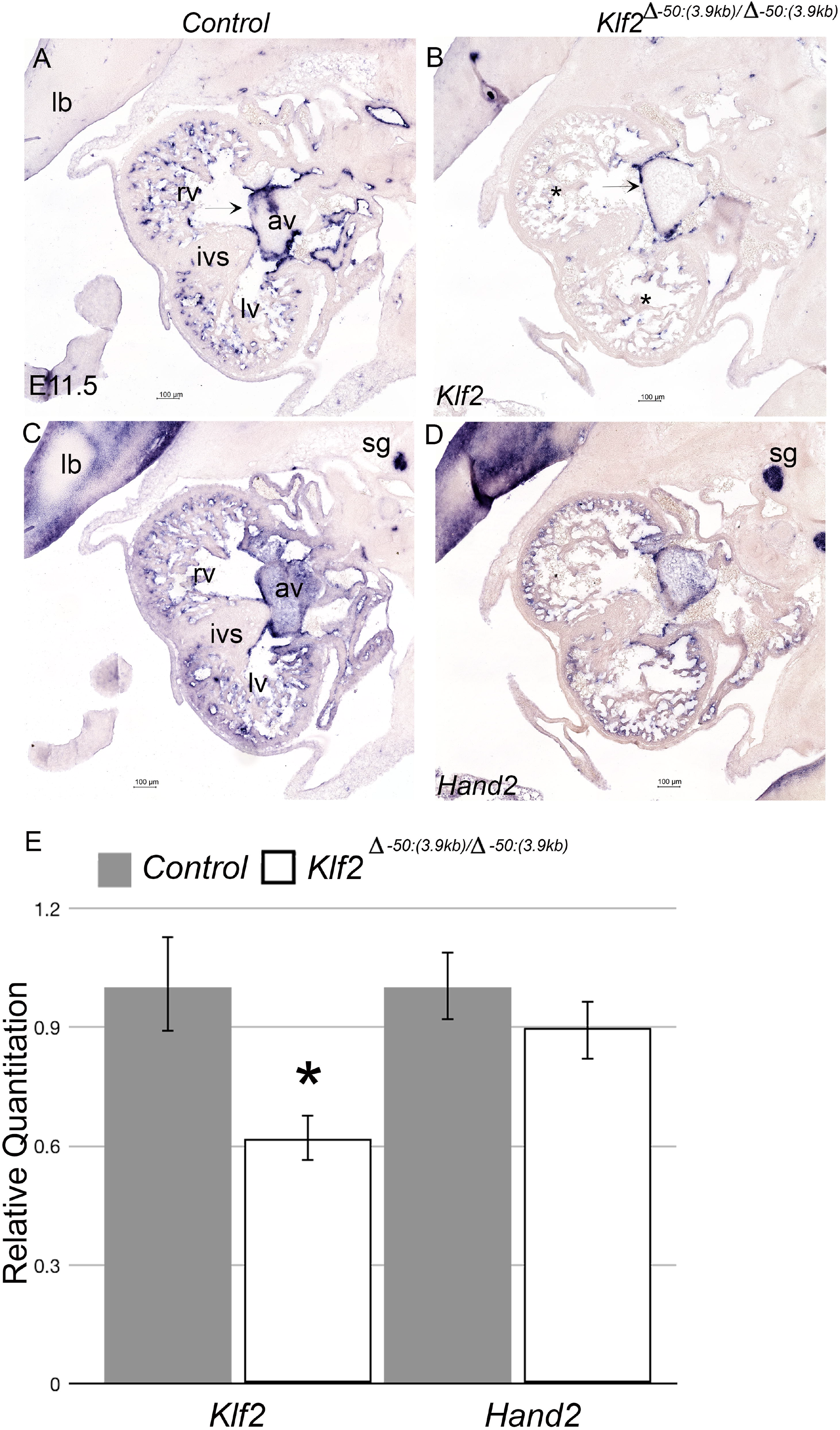
CRISPR/Cas9 mediated deletion of -50kb *Klf2* enhancer results in reduced *Klf2 ventricular endocardial* expression. **C. and E**. Section *in-situ* hybridization showing *Hand2* and *Klf2* expression in *Control* E11.5 embryos. **D. and F**. Section *in-situ* hybridization showing *Hand2* and *Klf2* expression in *Klf2*^*Δ-50kb(3*.*9kb)/Δ-50kb(3*.*9kb)*^ E11.5 embryos. Arrow indicates maintained expression within the endocardium overlying the cushions. Asterisk marks loss of gene expression within the ventricles of *Klf2*^*Δ-50kb(3*.*9kb)/Δ-50kb(3*.*9kb)*^ embryos. lb limb bud, rv right ventricle, lv left ventricle, ivs interventricular septum, sg sympathetic ganglia, av atrioventricular cushion.

## DISCUSSION

Loss of *Hand2* within the endocardium disrupts NOTCH signaling resulting in a hypotrabeculated single ventricle composed of hypervascularization free walls. (VanDusen, Casanovas, et al., 2014). To gain a better understanding on the gene regulatory networks in which HAND2 facilitates ventricular morphogenesis downstream of NOTCH1, we utilized single cell RNA-seq at E11.5, combined with established HAND2 DNA occupancy data (Laurent et al., 2017) to interrogate HAND2’s endocardial role in regulating endocardial gene regulatory networks. IPA analysis reveals a number of critical pathways known to be required for heart development that show misregulation within the identified endothelial/endocardial cell populations (Fig. 1). These analyses show disruption in several pathways, including wound healing, Pulmonary fibrosis and healing, tumor microenvironment (including HIF1*α* signaling) and most relevant to endocardial roles in cardiogenesis, Apelin pathway, which includes shear-stress response regulation (Fig. 1C, Supplemental Fig 4, Supplemental Spreadsheet 3). Collectively, these pathways contribute to endocardial response to vascularization of the myocardium, organ growth, and communication with the underlying myocardium coordinating septation and trabeculation. A number of significantly regulated genes as exampled by *Fn1, Ece1, and Edn1* exhibit altered expression, but do not exhibit robust HAND2 DNA occupancy in *cis*. Although such genes are influenced of HAND2 function, they are likely not directly transcriptionally regulated by HAND2, nevertheless their altered expression fits with HAND2 function in previous studies. In epicardial *Hand2* deletion, although *Fn1* expression is unaltered when comparing control and mutants, FN1 organization is altered within *H2CKO* epicardial cells (Barnes et al., 2011). During jaw morphogenesis, *Hand2* is established to lie downstream of EDN1 signaling and plays an important negative feedback role once activated, by repressing *Dlx5* and *Dlx6* expression within the ventral most portion of the mandible mesoderm (Barron et al., 2011; Charite et al., 2001; Clouthier et al., 2000; Vincentz et al., 2016).

By comparing highly misregulated genes with robust HAND2 DNA occupancy data located within CNS’s (Laurent et al., 2017), we chose five target genes, *Igf2, Igf2R, Ptn, Tmem108*, and *Klf2* to look for putative endocardial/endothelial HAND2-dependent enhancers (Fig. 3 and Supplemental Fig 5). CNS peaks bound to HAND2 from *Igf2, Tmem108*, and *Ptn* did not reveal any transcriptional activity (Supplemental Fig 5). More interestingly, we discovered three endocardial/endothelial enhancers, one CNS 5’ of the *Igf2R* TSS, and two CNS upstream of the *Klf2* TSS (Fig. 3).

The critical source of IGF2 for the heart is from the epicardium (Shen et al., 2015). Epicardial IGF2 diffuses into the heart where it can bind to its receptors, including IGF2R. Binding to IGF2R facilitates IGF2 degradation within the lysosomes (Harris & Westwood, 2012). Knockout of IGF2R within endothelial cells using *Tie2Cre* does not result in embryonic lethality; however, cardiac specific phenotypes have not been examined (Sandovici et al., 2022). Our data suggests that IGF2R plays a critical role within cardiac endothelium in a HAND2-dependent manner. We observe significant changes cell numbers within *H2CKO*s and *control*s (Supplemental Table 1) although it is yet to be determined if these changes are caused due to increased proliferation. Cell proliferation within E10.5 right ventricle of *Tie2Cre* mediated *H2CKO*s did not show significant differences (VanDusen, Casanovas, et al., 2014; Vandusen, Vincentz, et al., 2014)

Since KLF2 is a known master regulator of sheer-stress response and is a significant regulator within the Apelin regulatory network, we engineered a stable *Klf2* reporter line using the more robustly consistent endothelial/endocardial CNS located at -50kb of the *Klf2* TSS. Reporter expression analysis reveals that this *Klf2* CNS recapitulates all of *Klf2’s* endothelial/endocardial expression and is dependent on HAND2 only within the endocardium correlating directly with *Klf2* mRNA expression regulation (Fig. 2 and 4). KLF2 is expressed in regions of endothelium exposed to high sheer stress (Goddard et al., 2017). In the developing heart such high shear-stress regions include the endocardium overlying the developing valves and the developing ventricular trabeculae, with *Klf2* expression levels varying within regions of differential shear force (Goddard et al., 2017). Loss of HAND2 does not appear to impact *Klf2* expression within the regions of endocardium overlying the developing valve cushions where *Hand2* expression is already downregulated (VanDusen, Casanovas, et al., 2014; Vandusen, Vincentz, et al., 2014). Previous work characterizing conserved non-coding elements at *Klf2* genomic locus identified a 60bp enhancer element located 100bp upstream of *Klf2* transcriptional start site that is responsive to shear stress within mouse endothelial cells in culture (Huddleson et al., 2004). It is currently unclear if either the -23 or -50kb CNS enhancer are shear stress responsive but it is clear that the -50kb enhancer can recapitulate all *Klf2* mRNA expression domains during embryogenesis and the majority of its activity is HAND2-independent given HAND2 is not expressed within the systemic vasculature (Fig. 4). As one would expect, -50kb *Klf2* enhancer element contains other consensus binding sequences including Myocyte Enhancer Factor 2 (MEF2) family of transcription factors that are established regulators for vascular homeostasis and are transcriptional activators of *Klf2* (De Val & Black, 2009; Y. W. Lu et al., 2021). Analysis of DNA occupancy data shows that both -23kb and -50kb *Klf2* enhancers have conserved MEF2C binding (Akerberg et al., 2019). Given the established role of MEF2C endothelial integrity and homeostasis, it is not surprising that loss of *Hand2* does not lead to loss of vascular KLF2 expression.

Lineage tracing analysis shows that the endocardium is a primary of source of cells that eventually give rise to coronary vessels (Sharma, Chang, & Red-Horse, 2017). Studies in mouse models demonstrate that both endocardial and epicardial cells migrate into the myocardium to give rise to patent vessels (Sharma et al., 2017) and that these coronaries form within different zones of the myocardium, septum vs free walls of the ventricles (H. I. Chen et al., 2014). This suggests that coronary angiogenesis is driven by distinct mechanisms within different regions of the developing heart. The cellular origin of coronary vasculature is a source of some debate, the current consensus being that coronaries of the ventricular free wall are derived from the epicardium and the sinus venosus, whereas interventricular septal coronaries are derived from the ventricular endocardium (Phansalkar et al., 2021; Rhee et al., 2021; Zhang et al., 2016).

*Klf2* undergoes robust sheer stress response, as at least 50% of the highly flow regulated genes are dependent on *Klf2* upregulation (Parmar et al., 2006). *Klf2* expression within the endocardial cells of the ventricular wall fated to contribute to coronaries in these endocardial cells is HAND2-dependent. Indeed, one of the most striking observations in *H2CKO*s heart endocardium and vasculature is the persistent expression of *Lyve-1* beyond its normal endocardial downregulation by E13.5 (VanDusen, Casanovas, et al., 2014). LYVE1 expressing endocardium ultimately contribute to peripheral cardiac macrophages and developing lymphatic vasculature of the heart where vessel pressures are far less than encountered in blood vasculature (Pinto et al., 2012). It is an appealing idea that a defective shear-stress response of the ventricular endocardium could result in improper development/maturation of the ventricular endocardium into the correct sub-fates that result in hypervascularization of the ventricular walls composed of immature more lymphatic-like endothelium. Further support for this idea comes from multiple lines of evidence demonstrating that KLF2 inhibits angiogenesis by interacting with *VegfR2/Kdr* promoter (Bhattacharya et al., 2005), as loss of KLF2 also leads to hypervascularization (Kawanami et al., 2009). In our *H2CKO* data, we observe a modest increase in *Kdr* expression in cluster 7 (log2FC 0.13, not significant) which would be supportive of this possible mechanism.

Multiple genetic knockouts have been generated to study KLF2 function within endothelial cells. The *Klf2* systemic knockout is embryonically lethal between E12.5 to E14.5 due to severe intra-embryonic and intra-amniotic hemorrhaging (Kuo et al., 1997; Wani, Means, & Lingrel, 1998). Endothelial knockout of *Klf2* (and the related *Klf4*) using tamoxifen inducible *Cdh5-Ert2Cre* in 8- to 10-week adult mice causes vascular leakage leading to hemorrhaging and death (Sangwung et al., 2017). Endothelial knockout of *Klf2* using *Tie2Cre* exhibits increased systolic stroke volumes and high output heart failure leading to death at E14.5 due to abnormal vessel tone (Lee et al., 2006) and endocardial knockout of *Klf2* using *Nfatc1*^*Cre*^ results in embryonic lethality by E14.5 due to septal defects arising from a failure of cushion remodeling (Goddard et al., 2017). Moreover, work in zebrafish demonstrates that flow-responsive klf2 activates notch signaling, through a mechanism employing endocardial primary cilia (X. Li et al., 2020).

Given that *Klf2*^*Δ-50:(3*.*9kb)/Δ-50:(3*.*9kb*^ mice exhibit only a 40% reduction in *Klf2* endocardial expression and appears to maintain systemic vascular expression through other identified enhancers (Fig. 5), it is not surprising that that removal of this -50kb *Klf2* CNS does not result in embryonic lethality and that mice are viable and fertile. What we do not know currently, is the critical *Klf2* expression threshold that is causative to the observed embryonic vascular phenotypes or if any KLF2 endocardial-specific phenotypes contribute to the observed embryonic lethality. Collectively, these data demonstrate that HAND2 integrates endocardial transcriptional networks reaching beyond NOTCH pathway and that include sheer stress response revealing a number of important roles during endocardial morphogenesis.

## MATERIALS AND METHODS

### Mouse Strains and Genotyping

*Hand2*^*fx/fx*^ mice (Morikawa & Cserjesi, 2008)Jax strain 027727) and *Nfatc1*^*cre*^ (Wu et al., 2012) were genotyped as described previously (VanDusen, Casanovas, et al., 2014). The University of Michigan Transgenic Animal Model Core generated *LacZ* transgenic enhancer lines in the FVB background. 12 transmitting founder lines were screened for X-gal staining and enhancer activity. Transgenic founders and embryos were genotyped using primers spanning the enhancer and *HSP68* promoter 5’-AGCCTGTGAGAGAGACCCAT-3’ and 5’-GATGTTCCTGGAGCTCGGTA-3’. Genotyping for other alleles was carried out using Southern blots as previously described (George & Firulli, 2021). All animal maintenance and procedures were performed in accordance with the Indiana University School of Medicine protocol 20090, and University of Michigan School of Medicine. Animal work at Lawrence Berkeley National Laboratory (LBNL) was reviewed and approved by the LBNL Animal Welfare Committee.

### Single cell RNA-seq

E11.5 embryos were dissected in cold PBS and placed in PBS with 1% FBS solution on ice until dissociation (approximately 3 hours). Yolk-sac DNA was extracted (QuickExtract DNA Extraction Solution, Epicentre) and used for genotyping to distinguish heterozygous and homozygous *Hand2* conditional allele. The *Rosa*^*mTmG*^ allele fluorescence was used to determine *Nfatc1*^*cre*^ status. Dissected cardiac tissue was incubated in 750 μl TrypLE (ThermoFisher) for 5 min at 37^0^C, triturated with a 200-μl wide-bore pipette tip. The cell suspension was quenched with 750 μl DMEM with 10% FBS. Cells were filtered through a 30-μm cell strainer (MACS SmartStrainer), centrifuged at 300g for 5 min, and washed once with 750 μl PBS with 0.5% BSA. Cells were resuspended in 30 μl PBS with 0.5% BSA (10x Genomics). Single-cell droplet libraries from this suspension were generated using the Chromium NextGEM Single Cell 3’ Reagent Kits User Guide, CG000204 Rev D (10X Genomics, Inc), according to the manufacturer’s instructions. Briefly, each clean single cell suspension was counted with hemocytometer under microscope for cell number and cell viability. Only single cell suspensions with a viability of >90% and minimal cell debris and aggregation were used for further processing. The resulting library was sequenced in a custom program for 28b plus 91b paired-end sequencing on Illumina NovaSeq 6000. About 50K reads per cell were generated and 91% of the sequencing reads reached Q30 (99.9% base call accuracy).

Sequenced reads were aligned to a mouse transcriptome reference built from GRCm38.p6 (Genome Reference Consortium Mouse Build 38 patch release 6) combined with eGFP and dTomato gene sequences using the software 10x Genomics Cell Ranger 5.0.1 (Zheng et al., 2017). Reads from the cells associated with a total more than 1000 UMIs from Hemoglobin related genes (Hbb-bt, Hbb-bs, Hbb-bh2, Hbb-bh1, Hbb-y, Hba-x, Hba-a1, Hbq1b, Hba-a2 and Hbq1a) were excluded from further analysis. The downstream data exploration and differential gene expression analysis was conducted using the R package, Seurat V4 (Hao et al., 2021). As per the standard pre-processing workflow for scRNA-seq data in Seurat, cells with more than 2500 unique features were filtered out. The feature expression values for each cell were normalized using the standard “LogNormalize” method with default parameter values. The Seurat objects derived from WT and MT data were integrated using the anchors found using Canonical Correlation Analysis (CCA) with the neighbor search space specified using 1 to 20 dimensions (*FindIntegrationAnchors(****reduction****=“cca”, dims = 1:20)*). The integrated dataset was subjected to linear transformation followed by linear dimensionality reduction using Principal Component Analysis (PCA). Clusters were identified from the Shared Nearest Neighbor graph (*FindNeighbors(reduction = “pca”, dims = 1:20)*) with the resolution set to 0.5 (*FindClusters(resolution = 0*.*5)*) and were visualized using the Uniform Manifold Approximation and Projection (UMAP) non-linear dimensional reduction technique. For each cluster, the differentially expressed genes between *control* and *H2CKO* genotypes were called using a Wilcoxon Rank Sum test (*FindMarkers(****test*.*use****=“wilcox”)*). Genes with Bonferroni corrected p-values not more than 0.05 were considered significantly differentially expressed. For IPA analysis, the pathways relevant to the significantly differentially expressed genes (FDR <= 0.05) were identified using the Core Analysis of the IPA software (QIAGEN Inc., https://www.qiagenbio-informatics.com/products/ingenuity-pathway-analysis).

### CRISPR/Cas9 mediated deletion of -50kb *Klf2* enhancer

To generate the CRISPR-KO, single guide RNAs were designed flanking the -50kb *Klf2* enhancer by University of Michigan Transgenic Core 5’-CTACTACTTGGCAGGTTGGAGGG-3’ and 5’-GTCAAAGGGACCTGGTAGTTTGG-3’. Guide RNAs were tested for inducing chromosome breaks prior to microinjection. 114 potential founders were screened with PCR primers spanning the deletion, 5’-ATGTGTGTGCATCTGGGGAGCAGAG-3’ and 5’-CCAGAGTGACTTTTCAGGCACAGGGG-3’ which generates a 450bp product for the deleted allele. Primers within the deleted region were used to confirm a true indel, WT5’-CTTATAACCTCCATTTCCTCCTCTGGG-’3 and WT3’-CTTCGTGGTTTCCTGCTTGCTAAGATG-’3 that generates a 350bp product for the wildtype allele. PCR products from 31 positive founders were cloned and sequence verified to characterize the deletion. A probe for Southern blot was designed by using the following primers: 5’-CAAGGCCTTCCAGTACCAGG-3’ and 5’-TCTCAGTGGAGCTTGCTGTG-3’ to clone out a 332bp fragment from murine genomic DNA. The probe detects an RFLP in EcoRV digested genomic DNA, 9.5kb in wildtype allele and 5.6kb in the CRISPR deleted allele (*Klf2* ^*Δ-50kb(3*.*9kb)*^). Selected founders were outcrossed for two generations before being bred to homozygosity.

### Transgenic mouse reporter assays

Mouse transgenesis at LBNL was performed in *Mus musculus* FVB strain mice. Animals of both sexes were used in these analyses and mouse embryos were excluded from further analysis if they did not encode the reporter transgene or if the developmental stage was not correct. For validation of *in vivo* enhancer activities, enSERT was used for site-directed insertion of transgenic constructs at the H11 safe-harbor locus (Kvon et al. 2020). This method is based on pronuclear co-injection of *Cas9*, sgRNAs and a H11-homology arms-containing targeting vector encoding a candidate enhancer element upstream of a minimal promoter and a reporter protein (Kvon et al. 2020, Osterwalder et al. 2022). Related genomic enhancer coordinates are listed in Table 1. Predicted enhancer regions were PCR-amplified from mouse genomic DNA from wildtype FVB mice and cloned into a modified targeting vector encoding a human *beta-globin* minimal promoter upstream of a LacZ reporter. Embryos were excluded from further analysis if they did not contain a reporter transgene. CD-1 females served as pseudo-pregnant recipients for embryo transfer to produce transgenic embryos which were collected at E11.5 and stained with X-gal using standard techniques (Kothary 1989, Osterwalder et al. 2022). For generation of stable enhancer-reporter lines, a *Hsp68-LacZ* reporter-based vector for random integration was used (Vincentz et al., 2012). Embryos were harvested from timed mating’s at the timepoints indicated and pre-fixed in 2% paraformaldehyde-0.2% glutaraldehyde and stained as previously described (VanDusen, Casanovas, et al., 2014; Vincentz et al., 2019). After overnight staining at room temperature, embryos were post-fixed in 4% paraformaldehyde prior to imaging and sectioning.

### Histology

If stained, LacZ stained embryos were post-fixed, washed in PBS, dehydrated, embedded, sectioned, and Nuclear Fast Red (NFR) stained as previously described (George & Firulli, 2021; Vincentz et al., 2019). Images were acquired on the Keyence BZ-X800 florescence microscope system or the Leica DM5000 B compound microscope.

### Cloning

Conserved non-coding putative HAND2 binding regions were cloned out from genomic mouse DNA using the following primers: *Igf2* 5’-GAGAAGCTGGCAGATCAGGCTGTG-3’ and 5’-TGCTTCTGTTGAGAGGAGACAGTCTGG-3’, *Igf2r* 5’-TTGCCTGCATGTAAGTGTGCCTGG-3’ and 5’-TGTCTCTCAGGCTTCCTGTCTGGC-3’, *Ptn* 5’-ATTTCAGCTGGACTGCCATGGCAG-3’ and 5’-GGCTGGAAGAGGAGGCAAACAGAG-3’, *Tmem108* 5’-CATCATCACCATCACCATCGTCGTCG-3’ and 5’-GTATGCAGTGGACCTCTTTGACTTGTCAG-3’, *Klf2* enhancer -23kb element 5’-ATCTGTCCACCTCTACCTTCCA-3’ and 5’-AGTGGCTCTGACAACCTGAGAT-3’, *Klf2* enhancer -50kb element 5’-TGAACCTCCATTGATACACACC-3’ and 5’-GTCCCTAAGGATCATGTTGAGC-3’. Amplified sequences were Gibson (NEB) cloned into the *pCR4-bG::lacZ-H11* enSERT vector and used to generate F0 enhancer transgenics. Briefly, the enSERT system uses CRISPR/Cas9 mediated site directed transgenesis at the murine H11 locus resulting in genomic integration of the human *beta-globin* promoter with the enhancer element to be tested and the *lacZ* reporter cassette (Kvon et al., 2020). F0’s are harvested at E11.5 for *lacZ* staining and analysis.

To generate the -50kb *Klf2* stable transgenic allele, primers corresponding to genomic region chr8:74791237-74793083 (mm9) were used 5’-AAGGGCCAGATGTGCTGAAA-3’ and 5’-GGCTGGTCTCGAACTCACAA-3’, and cloned into *HSP68-LacZ* vector backbone as described previously (Vincentz et al., 2019) and used to create stable *β-gal* expressing mouse transgenic lines.

### *In-situ* hybridization

Section in situ hybridizations (ISH) were performed on 10-μm paraffin sections as described previously (George & Firulli, 2021). Whole mount in situ hybridizations were performed using E10.5 day embryos as described previously (George & Firulli, 2021). Antisense digoxygenin-labeled riboprobes were synthesized using T7, T3, or SP6 polymerases (Promega) and DIG-Labeling Mix (Roche) using the following plasmid templates: *Hand2, Klf2*.

### Quantitative real time PCR

Total RNA was isolated from E11.5 ventricles using the High Pure RNA Isolation Kit (Roche). RNA was used to synthesize cDNA using the High-Capacity cDNA Reverse Transcription Kit (Applied Biosystems). For qRT-PCR, cDNA was amplified using TaqMan Probe-Based Gene Expression Assays (Applied Biosystems) to quantify gene expression. qRT-PCR reactions were run on the QuantStudio 3 Real-Time PCR System (ThermoFisher). Normalization to Glyceraldehyde 3-phosphate dehydrogenase (GAPDH) was used to determine relative gene expression and statistical analysis was automatically applied by the instrument software. Significance of qRT-PCR results were determined by a two-tailed student’s t-test followed by post hoc Benjamini-Hochberg FDR correction as automatically calculated by the QuantStudio3 qRT-PCR thermal cycler software analysis package. Data are presented as Relative Quantitation values where error bars depict the maximum and minimum values of each series of samples. A minimum n of 8 is used in all assays.

### ChIP PCR assays

For ChIP assays, NIH3T3 cells were transfected with Lipofectamine3000 with plus reagent (Invitrogen) according to manufacturer’s instructions with pCS2+Myc-Hand2, pCS2+Myc-E12, or pCS2 control constructs as indicated. After culturing for 48 hours, SimpleChIP plus enzymatic chromatin IP kit (Cell Signaling Technologies) was used for ChIP experiment as per manufacturer recommendations and PCR was used to detect ChIP products run out on agarose gel.

## ACKNOWLEDGEMENTS

We thank Danny Carney and Chloe Ferguson for technical assistance. *Klf2* expression vector used to generate ISH probe was a kind gift from Jonathan A Epstein. We thank Nathan VanDusen for helpful comments.

## Author Contributions

RMG and B.F. designed and performed experiments, wrote, and edited the manuscript; RP and DBR Bioinformatic analysis, BJM, LP and MO transgenic mouse construction and analysis. A.B.F. designed and performed experiments, performed data interpretation, wrote, and edited the manuscript.

## Data availability

This manuscript contains sequence data is deposited on GEO - GSE210221. We agree to make all mice engineered by us and all data freely available.

## Supplemental Legends

**Supplemental Spreadsheet 1: Assigning cluster identity to *H2CKO*s and *control* using conserved markers**

Comparing *H2CKO*s and *control* gene expression in the cluster against that in other clusters to determine identity. Genes of interest are highlighted (matching the color of the respective cluster in Fig 1A, Cluster 0 red, Cluster 1 pink, Cluster 2 brown, Cluster 3 orange, Cluster 4 light purple, Cluster 5 dark purple, Cluster 6 light green, Cluster 7 light blue, Cluster 8 dark green, Cluster 9 dark blue, Cluster 10 grey, Cluster 11 yellow, Cluster 12 black). Cluster_ID identify individual clusters. Gene and ID (ensembl) identify the gene. WT_avg_log2FC indicates Log2 fold change for WT calculated by taking average WT gene expression in cluster divided by average WT gene expression in all other clusters. WT_pct1 indicates percent of WT cells in the cluster with gene being expressed. WT_pct2 indicates percent of WT cells in all other clusters with gene being expressed. WT_p_val_adj indicates adjusted p-value for WT (with Bonferroni correction), and description.

**Supplemental Spreadsheet 2: Differentially expressed genes in *H2CKO*s vs *control***

Column WT_avg_log2FC indicates log2 fold change for *control* (Average WT gene expression in cluster divided by average WT gene expression in all other clusters). Column WT_pct.1 indicates percent of *control* cells in the cluster with gene being expressed. Column WT_pct.2 indicates percent of *control* cells in all other clusters with gene being expressed. Column WT_p_val_adj indicates adjusted p-value for *control* (with Bonferroni correction). WT_x(cluster number) indicates the average gene expression for each gene in log2 space in *control*. MT_x(cluster number) indicates the average gene expression for each gene in log2 space in *H2CKO*s. Endocardial *Hand2* (in clusters 7 and 9) highlighted in red, genes highlighted in yellow were selected for F0 transgenic embryo analysis, and genes of interest are highlighted in blue.

**Supplemental Spreadsheet 3: Canonical pathways significantly changed in cluster 7 in *H2CKO*s vs *Control***.

List of the most significant canonical pathways across the entire dataset. For each pathway, the significance (−log(p-value)) is calculated by right-tailed Fisher’s exact test. The significance indicates the probability of association of molecules from the dataset with the canonical pathway by random chance alone.

**Supplemental Table 1: Differential abundance analysis on *H2CKO*s and *control***. Number of barcodes represent values of unique molecular identifiers corrected for multiples and *Hbb* contamination. Adjusted p-value is calculated by using Fishers p-value followed by adjustment for multiple testing using Benjamini-Hochberg (BH) method. Description of cell numbers indicates clusters that have significant changes in barcodes between *H2CKO*s and *control*.

CM cardiomyocytes, cNCC cardiac neural crest cells, RBC red blood cells, OFT outflow tract mesenchyme, EndoMT endothelial to mesenchymal transition.

Supplemental Table 2: List of downstream KLF2 targets that significantly changed in H2CKO endocardium.

**Supplemental Figure 1: Cluster identity analysis**

UMAP data showing gene expression in each of the indicated clusters based on data contained within Supplemental Spreadsheet 1.

**A**. Gene expression in individual clusters. Scale bar showing Log2FC values for each gene

**B**. UMAP plot of all barcodes (11,640) captured by scRNA-seq of *control* (*Hand2*^*fx/+*^) and *H2CKO* (*Nfatc1*^*cre*^*Hand2*^*fx/fx*^) E11.5 hearts. Cluster identification by comparing gene expression in individual *control* clusters comparing each cluster to all others combined.

**Supplemental Figure 2: *Hand2* expression in *H2CKO* and *Control* clusters**

Violin plots showing *Hand2* expression across cluster. Orange plots represent *H2CKO* and green plots represent *control*. The line in the middle of the box plot is median while the edges are the first and the third quartiles respectively. The vertical line represents the complete range of the expression (min to max).

**Supplemental Figure 3: IPA on differentially expressed genes from *H2CKO*s cardiomyocytes**

Graphical summary of differentially expressed genes from cluster 0 (cardiomyocytes).

**Supplemental Figure 4: IPA on differentially expressed genes from *H2CKO*s endocardium**

Graphical summary of differentially expressed genes from cluster 7 (endocardial cells).

**Supplemental Figure 5: F0 reporter expression analysis of target genes showing both altered gene expression and HAND2 DNA-occupancy**

**A**. *Igf2* genomic locus showing conserved non-coding regions (green solid boxes), transcriptional start site (+1), relative location of enhancer element (+70kb, red outline). HAND2^3xFlag^ ChIP-seq data (Laurent et al., 2017) showing genomic regions of HAND2 binding.

**B. B’**. +70kb HAND2 binding conserved non-coding region at *Igf2* locus used to make transgenic F0 embryos and *lacZ* staining results. Representative whole mount image of E11.5 transgenic embryo. Numbers of transgenic F0 embryos obtained, 9. Numbers of F0 embryos that showed staining, 0.

**C**. *Tmem108* genomic locus showing conserved non-coding regions (green solid boxes), transcriptional start site (+1), relative location of enhancer element (red outline). HAND2^3xFlag^ ChIP-seq data (Laurent et al., 2017) showing genomic regions of HAND2 binding.

**D, D’**. HAND2 binding conserved non-coding region at *Tmem108* locus used to make transgenic F0 embryos and *lacZ* staining results. Representative whole mount image of E11.5 transgenic embryo. Numbers of transgenic F0 embryos obtained, 3. Numbers of F0 embryos that showed staining, 0.

**E**. *Ptn* genomic locus showing conserved non-coding regions (green solid boxes), transcriptional start site (+1), relative location of enhancer element (+67kb, red outline). HAND2^3xFlag^ ChIP-seq data (Laurent et al., 2017) showing genomic regions of HAND2 binding. **F, F’** +67kb HAND2 binding conserved non-coding region at *Ptn* locus used to make transgenic F0 embryos and *lacZ* staining results. Representative whole mount image of E11.5 transgenic embryo. Numbers of transgenic F0 embryos obtained, 3. Numbers of F0 embryos that showed endocardial staining, 0.

**Supplemental Figure 6: -21kb *Igf2r* enhancer CLUSTWAL alignment**

Evolutionary conservation within -21kb *Igf2r* enhancer. Black boxed sequences indicate E/D boxes.

**Supplemental Figure 7: *LacZ* stable transgenic lines of -50kb *Klf2* enhancer** Representative images of stable -50kb *Klf2* conserved non-coding element *lacZ* transgenic lines at E11.5. Scale bar 1mm.

**Supplemental Figure 8: -50kb *Klf2* enhancer CLUSTWAL alignment**

Base pairs in light blue show regions of evolutionary conservation within -50kb *Klf2* enhancer. Black boxed sequences indicate E/D boxes. Ebox 1, 2, and 3 assayed by ChIP are highlighted in green and marked.

**Supplemental Figure 9: Design and validation of the *Klf2***^*Δ****-50kb(3*.*9kb)***^ **gene edited allele**

**A**. Deletion of -50kb *Klf2* enhancer by CRISPR/Cas9 with 5’- and 3’-guide RNA (gRNA, red scissors). Total size of deletion is 4300bp exceeds the size of the identified enhancer and was necessary to find efficacious gRNAs. PCR primers used for genotyping (in blue), wildtype product size 450bp, deletion product size 350bp. *EcorV* restriction digest (blue triangles) are used to generate RFLPs for Southern blots.

**B**. Southern blots for the wildtype (*Klf2*^*wt/wt*^) RFLP migrate at 9.5kb and successful gene-edited enhancer deletion mutants (*Klf2*^*Δ-50kb(3*.*9kb)/Δ-50kb(3*.*9kb)*^) RFLPs migrate at 5.5kb. Heterozygous alleles (*Klf2* ^*Δ-50kb(3*.*9kb)/wt*^) contain both RFLPs.

