## Supplemental Tables and figures for "Single cell evaluation of endocardial HAND2 gene regulatory networks reveals critical HAND2 dependent pathways impacting cardiac morphogenesis"

**Supplemental Table 1:**

| KLF2 transcriptional activity on target gene | Target Gene | Significantly regulated in *H2CKO* endocardium | Function related to KLF2 | HAND2 DNA occupancy? |
|---|---|---|---|---|
| Repressor | Tie2 (TEK) | Significantly down in cluster 7 and 9 | Angiogenesis | No |
| Activator | Hif1a | Significantly down in cluster 7 and 9 | Angiogenesis | No |
| Activator | SMAD4 | Significantly down in cluster 7 and 9 | Inflammation | No |
| Activator | SMAD7 | Significantly down in cluster 7 | Inflammation | No |
| Repressor | MAPK (MAPK1) | Down in cluster 7 (significant) and 9 | Oxidative stress | No |
| Activator | IL6 | Significantly down in cluster 7 and 9 | Anti-thrombosis | No |
| Activator | Cav1 | Up in cluster 7 (significant) and 9 | Vascular tone | No |

**Supplemental Table 1: List of downstream KLF2 targets that are significantly changed in *H2CKO* endocardium.**

**Supplemental Table 2**

| Cluster | Cluster Identity | *Control* barcode count within the cluster | *H2CKO* barcode count within the cluster | Adjusted p-value (BH) | Log 2 Odds Ratio | Description of cell numbers |
|---|---|---|---|---|---|---|
| **0** | CM | 1,067 | 1,731 | 4.77E-16 | -0.65 | Significant increase in *H2CKO*s |
| **1** | CM | 784 | 1,588 | 4.77E-16 | -1.01 | Significant increase in *H2CKO*s |
| **2** | EndoMT | 652 | 378 | 4.77E-16 | 1.09 | Significant decrease in *H2CKO*s |
| **3** | OFT CM | 468 | 526 | 6.90E-01 | 0.04 | Not significant |
| **4** | cNCC | 440 | 300 | 4.34E-13 | 0.81 | Significant decrease in *H2CKO*s |
| **5** | cNCC | 416 | 270 | 3.42E-14 | 0.88 | Significant decrease in *H2CKO*s |
| **6** | Fibroblasts | 454 | 140 | 4.77E-16 | 2.00 | Significant decrease in *H2CKO*s |
| **7** | Endocardium | 132 | 432 | 4.77E-16 | -1.57 | Significant increase in *H2CKO*s |
| **8** | Epicardium | 412 | 130 | 4.77E-16 | 1.95 | Significant decrease in *H2CKO*s |
| **9** | Endocardium | 235 | 227 | 6.72E-02 | 0.27 | Not significant |
| **10** | RBCs | 152 | 216 | 6.41E-02 | -0.31 | Not significant |
| **11** | Conduction CM | 136 | 203 | 2.53E-02 | -0.38 | Not significant |
| **12** | Leukocytes | 60 | 91 | 1.09E-01 | -0.40 | Not significant |

**Supplemental Table 2: Differential abundance analysis on *H2CKO*s and *Controls*.**
Number of barcodes represent values of unique molecular identifiers corrected for multiples and *Hbb* contamination. Adjusted p-value is calculated by using Fishers p-value followed by adjustment for multiple testing using Benjamini-Hochberg (BH) method. Description of cell numbers indicates clusters that have significant changes in barcodes between *H2CKO*s and *Controls*.
CM cardiomyocytes, cNCC cardiac neural crest cells, RBC red blood cells, OFT outflow tract mesenchyme, EndoMT endothelial to mesenchymal transition.

**Supplemental Figure 1: Cluster identity analysis**

**A**

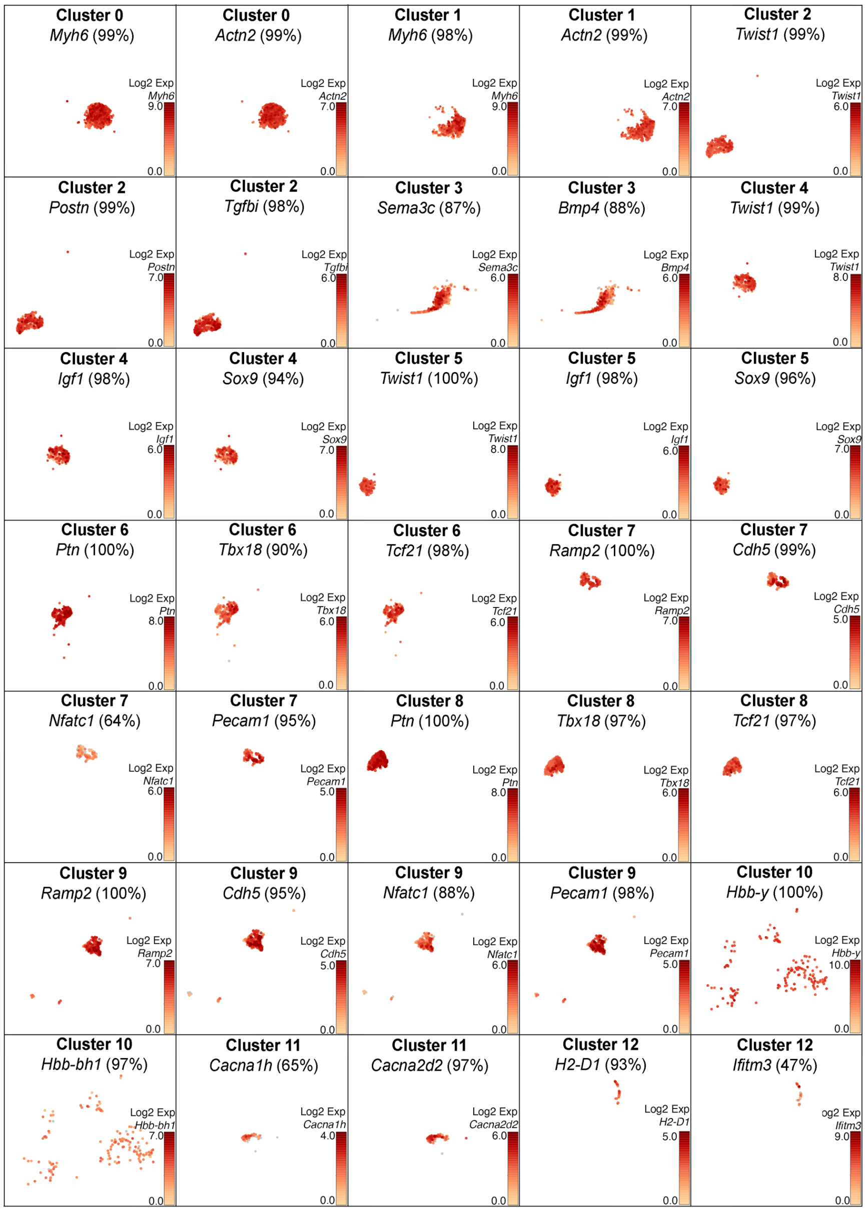


**B**

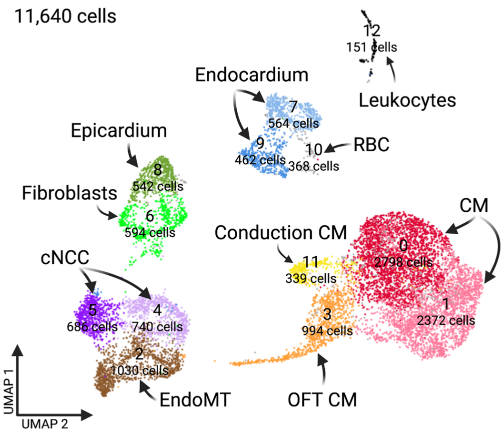

**Supplemental Figure 1: Cluster identity analysis**

UMAP data showing gene expression in each of the indicated clusters based on data contained within Supplemental Spreadsheet 1.

**A**. Gene expression in individual clusters. Scale bar showing Log2FC values for each gene

**B**. UMAP plot of all barcodes (11,640) captured by scRNA-seq of *control* (*Hand2*$^{fx/+}$) and *H2CKO* (*Nfatc1*$^{cre}$*Hand2*$^{fx/fx}$) E11.5 hearts. Cluster identification by comparing gene expression in individual *control* clusters comparing each cluster to all others combined.

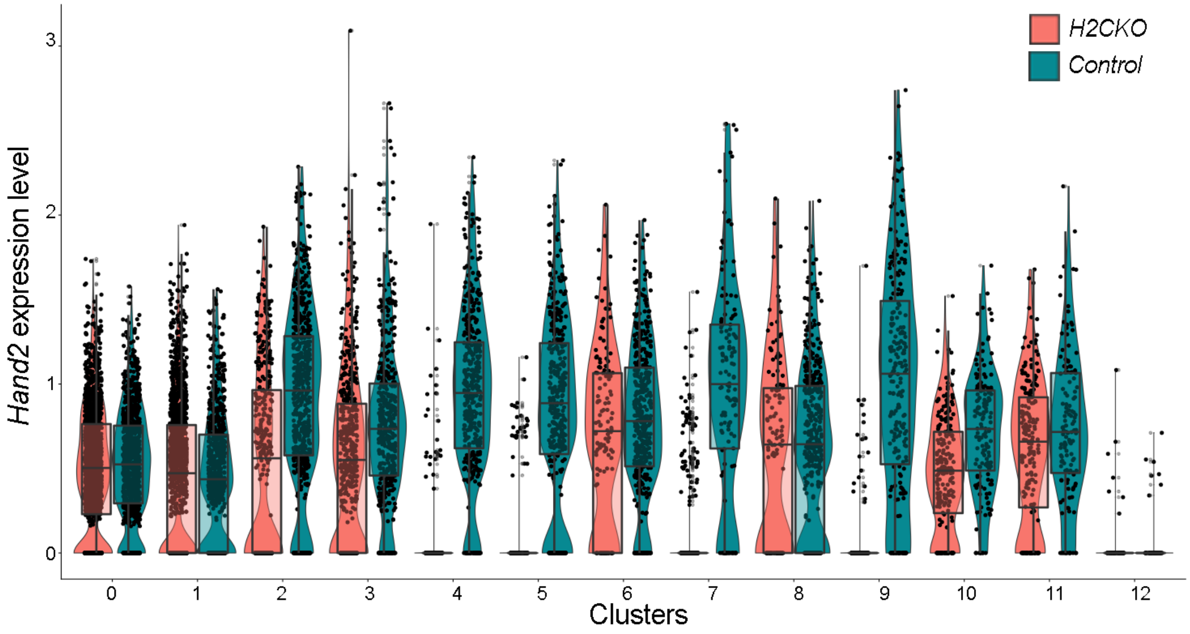


**Supplemental Figure 2: *Hand2* expression in *H2CKO* and *Control* clusters**

Violin plots showing *Hand2* expression across cluster. Orange plots represent *H2CKO* and green plots represent *control*. The line in the middle of the box plot is median while the edges are the first and the third quartiles respectively. The vertical line represents the complete range of the expression (min to max).

**Supplemental Figure 3:**
**Graphical summary of differentially expressed genes from cluster 0 (cardiomyocytes).**

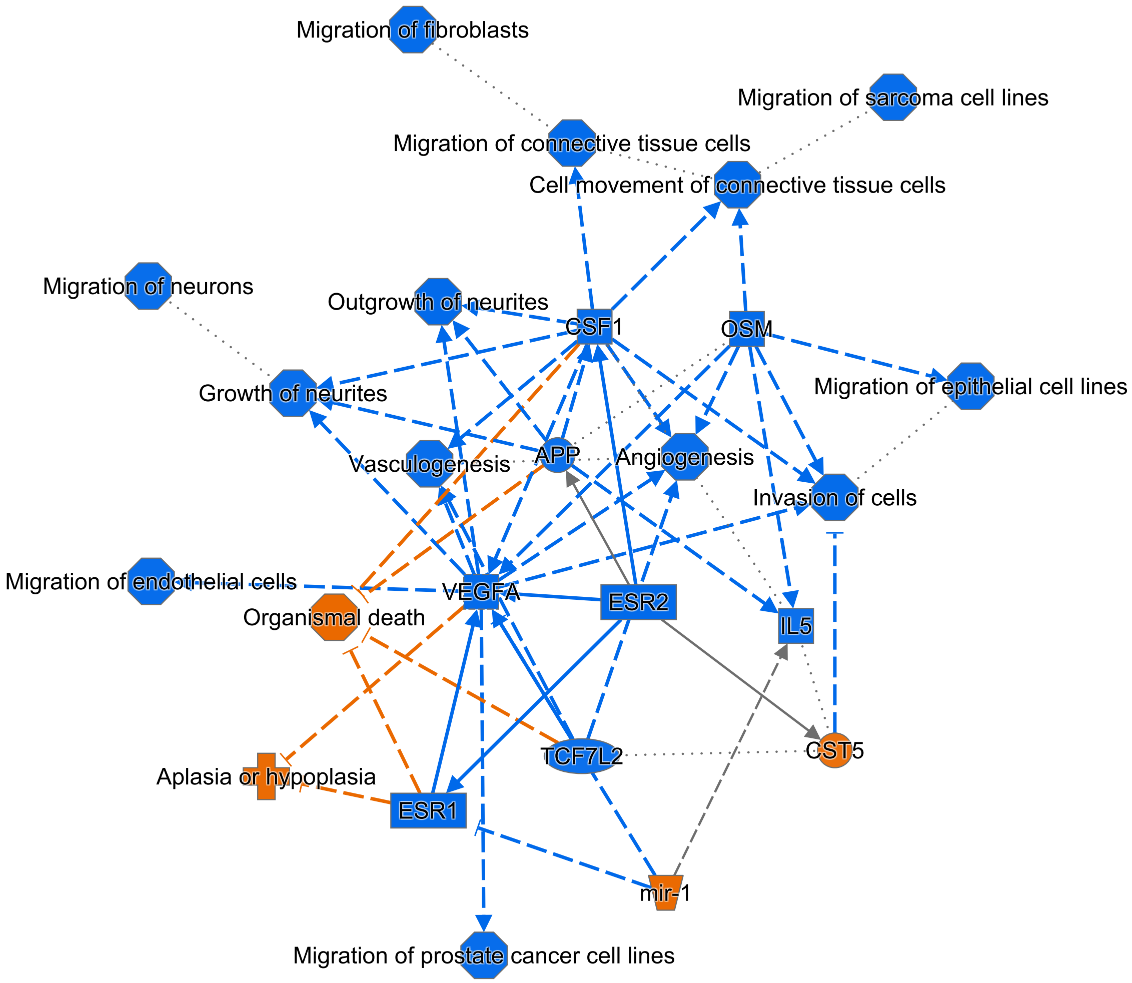


© 2000-2022 QIAGEN. All rights reserved.

**Supplemental Figure 3: IPA on differentially expressed genes from *H2CKOs* cardiomyocytes**

Graphical summary of differentially expressed genes from cluster 0 (cardiomyocytes).

**Supplemental Figure 4:**
**Graphical summary of differentially expressed genes from cluster 7 (endocardial cells).**

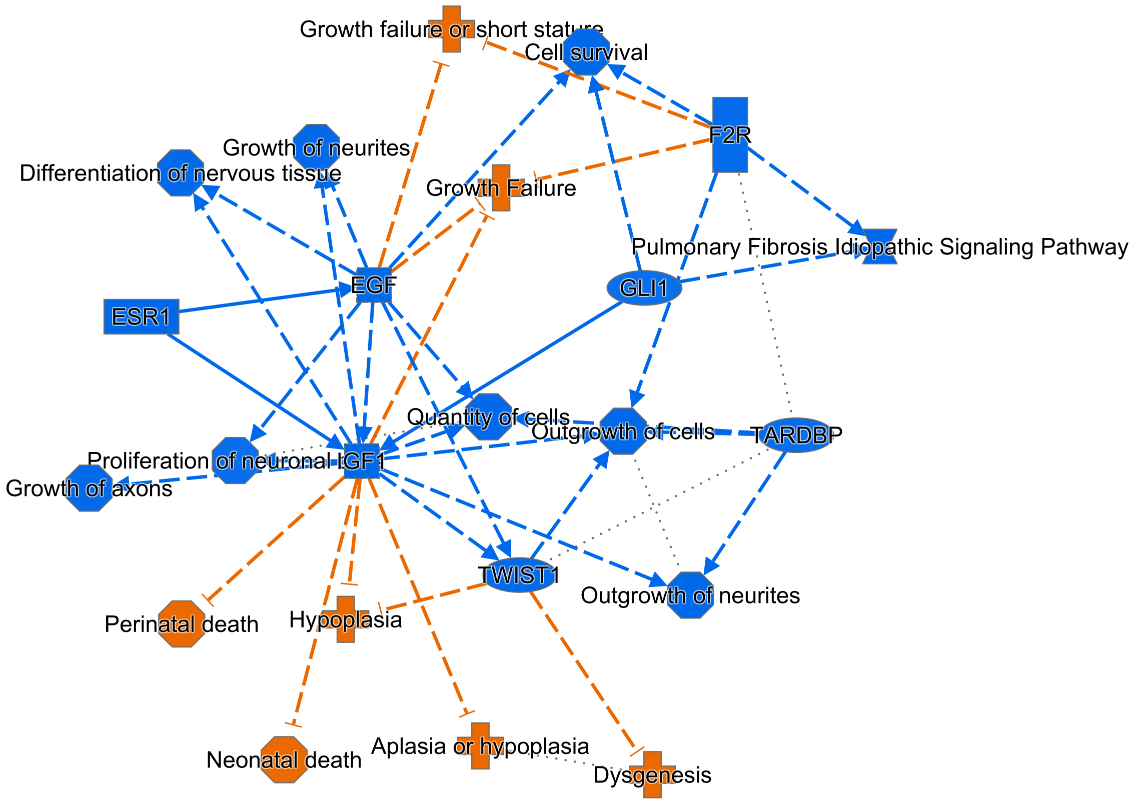


© 2000-2021 QIAGEN. All rights reserved.

**Supplemental Figure 4: IPA on differentially expressed genes from *H2CKO*s endocardium**

Graphical summary of differentially expressed genes from cluster 7 (endocardial cells).

### Supplemental Figure 5

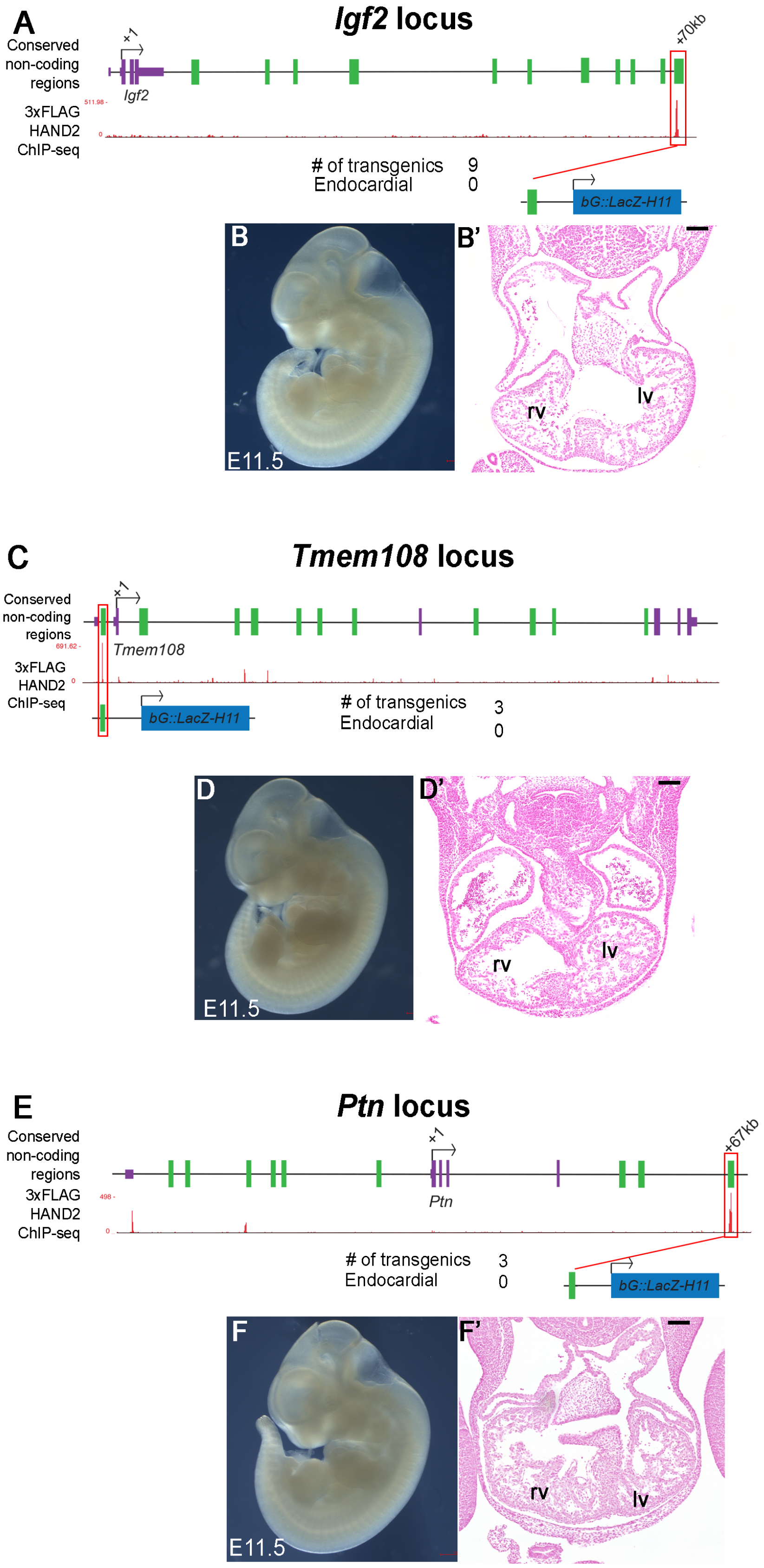

**Supplemental Figure 5: F0 reporter expression analysis of target genes showing both altered gene expression and HAND2 DNA-occupancy**

**A.** *Igf2* genomic locus showing conserved non-coding regions (green solid boxes), transcriptional start site (+1), relative location of enhancer element (+70kb, red outline). HAND2$^{3xFlag}$ ChIP-seq data (Laurent et al., 2017) showing genomic regions of HAND2 binding.

**B. B'.** +70kb HAND2 binding conserved non-coding region at *Igf2* locus used to make transgenic F0 embryos and *lacZ* staining results. Representative whole mount image of E11.5 transgenic embryo. Numbers of transgenic F0 embryos obtained, 9. Numbers of F0 embryos that showed staining, 0.

**C.** *Tmem108* genomic locus showing conserved non-coding regions (green solid boxes), transcriptional start site (+1), relative location of enhancer element (red outline). HAND2$^{3xFlag}$ ChIP-seq data (Laurent et al., 2017) showing genomic regions of HAND2 binding.

**D, D'.** HAND2 binding conserved non-coding region at *Tmem108* locus used to make transgenic F0 embryos and *lacZ* staining results. Representative whole mount image of E11.5 transgenic embryo. Numbers of transgenic F0 embryos obtained, 3. Numbers of F0 embryos that showed staining, 0.

**E.** *Ptn* genomic locus showing conserved non-coding regions (green solid boxes), transcriptional start site (+1), relative location of enhancer element (+67kb, red outline). HAND2$^{3xFlag}$ ChIP-seq data (Laurent et al., 2017) showing genomic regions of HAND2 binding.

**F, F'** +67kb HAND2 binding conserved non-coding region at *Ptn* locus used to make transgenic F0 embryos and *lacZ* staining results. Representative whole mount image of E11.5 transgenic embryo. Numbers of transgenic F0 embryos obtained, 3. Numbers of F0 embryos that showed endocardial staining, 0.

```
Mus_musculus        CCCCTTTTCATATTCACATGATTTTTTTTTT-CCTTTGGGTGGTATGGATGTTTTGCCTGCATGTAAGTGTGCCTGGGACTGGAAGAGGCCAGAAGATGCCCCC-GGAATTGGAGCTACA
Mus_spretus         CCCCTTTTCATATTCACATGATTTTTTTT-T-CCTTTGGGTGGTATGGATGTTTTGCCTGCATGTAAGTGTGCCTGGGACTGGAAGAGGCCAGAAGATGCCCCCAGGAATTGGAGCTACA
Mus_caroli          CCCCTTCTCATATTCACATGATTTTTTTTTCCCCTTTGGGTGGTATGGATGTTTTGCCTGCATGTAAGTGTGCCCGGGACTGGAAGAGGCCAGAAGATGCCCCC-GGAATTGGAGCTACA

Mus_musculus        AACAGTTGTGAGCTGCCATGTGGGTACTGGGAATCAAACCCAGGCCCTCCCAAGGAG---CCCCCAAGCACTCTTGACCACTGAACAATCTCTAGCCCCTGAAACACTTGTTTCTAAGAA
Mus_spretus         AACAGTTGTGAGCTGCCATGTGGGTACTGGGAATCAAACCCAGGCCCTCCCAAGGAG---CCCCCAAGCACTCTTGACCACTGAGCAATCTCTAGCCCCTGAAACACTTGTTTCTAAGAA
Mus_caroli          GACAGTTGTGAGCTGCCATGTGGGTACTGGGAATCAAACTCAGGCCCTCCCAAGGAGCAGCACCCAAGCACTCTTGACCACTGAGCAAGCTCTAGCCCCTGAAACACTTGTTTCTAAGAA

Mus_musculus        TGGATCAGTTGTCACAGAGGTTGACTCATGCAAGCTTAGGTCAGCCTCATTCTATTTTGGCATGGCAGAAATGCCAGCTCAGGCACAGAAACTCCCCTGGCAGATCTGGGGACAGTGGGT
Mus_spretus         TGGATCAGTTGTCACAGAGGTTGACTCATGCAAGCTTAGGTCAGCCTCATTCTATTTTGGCATGGCAGAAATGCCAGCTCAGGCACAGAAACTCCCCTGGCAGACCTGGGGACAGTGGGT
Mus_caroli          TGGACCAGTTGTCACAGAGGTTGACTCATGCAAGCTTAGGTCAGCCTCATTCTATTTTGGCATGGCAGAAATGCCAGCTCAGGCACAGAAACTCCCCTGGCAGACCTGGGGACAGTGAGT

Mus_musculus        TTATGGGCTCCCATCCCTGCTTCCTGTCCACCCTTTGACTTTGGACTGGTTGTTTAATCTCCGTTCCGCTTTCTTCATCCAGAGCAGAGGTCAGCAAACTCATCAAAGGGTCAGGCCGTA
Mus_spretus         ATATGGACTCCCATCCCTGCTTCCTGTCCACCCTTTGACTTTGGACTGGTTGTTTAATCTCCGTTCCGCTTTCTTCATCCAGAGCAGAGGTCAGCAAACTCATCAAAGGGTCAGGCCGTA
Mus_caroli          ATATGGGCTCCCATCCCTGCTTCCTGTCCACCCTTTGACTTTGGACTGGTTGTTTAATCTCCGTTCCACTTTCTTCATCCAGAGCAGAGGTCAGCAAACTCATCAAAGGGTCAGGCCGTA

Mus_musculus        AACTGTTTAAGCTGTATGGATCCTATGGCTTTCCGGCACAGCTACTCAGCTCTGCCCCTGTGGCATGAGAACGGCCTCAGGCAACGTGTGAGCCTATGTACGTGGCAGTGAGCCAAGAAA
Mus_spretus         AACTGTTTAAGCTGTATGGATCCTATGGCTTTCCGGCACAGCTACTCAGCTCTGCCCCTGTGGCATGAGAACGGCCTCAGGCAACGTGTGAGCCTATGTACATGGCAGTGAGCCAAGAAA
Mus_caroli          AACTGTTTAAGCTGTATGGATCCTATGGCTTTCCGGCACAGCTACTCAGCTCTGCCCCTGTGGCATGAGAACGGCCTCAGGCAACGTGTGAGCCTATGTACGTGGCAGTGAGCCAAGAAA

Mus_musculus        GCTTCACTTTTGAAAATAGCCAGATATGGTTCCATGTCTATAGCTTGCCAGCCCCTGATCAGATGACGCCCCCCTCCCCAGCCCCCTGCAAATGGAAGAATGTGGAAGACTTTACGGTTG
Mus_spretus         GCTTCACTTTTGAAAATAGCCAGATATGGTTCCATGTCTATAGCTTGCCAGCCCCTGATCAGATGACGCCCCCCTCCCCAGCCCCCTGCAAATGGAAGAATGTGGAAGACTTTACGGTTG
Mus_caroli          GCTTCACTTTTGAAAATAGCCAGATATGGTTCCATGTCTATAGCTTGCCAGCCCCTGATCAGATGACGCCCCCCTCCCCAGCCCCCTGCAAATGGAAGAATGTGGAGGACTTTACGGTTG

Mus_musculus        ACTGTTCAGTGTCTGTCACACTCCAGCTGATAGTGTAAGCCAAACAAACATGTCCATCCCCCGGGTACACTTAAGCCTGTGTGTGAAAGATGTAAGGAGATGATTACCAGAGCTCCTGGG
Mus_spretus         ACTGTTCAGTGTCTGTCACACTCCAGCTGATAGTGTAAGCCAAACAAACATGTCCATCCCCCGGGTACACTTAAGCCTGTGTGTGAAAGATGTAAGGAGATGATTACCAGAGCTCCTGGG
Mus_caroli          ACTGTTCAGTGTCTGTCACACTCCAGCTGATAGTGTAAGCCAAACAAACATGTCCATCCCCCGGGTACACTTAAGCCTGTGTGTGAAAGATGTAAGGAGATGATTACCAGAACTCCTGGG

Mus_musculus        AAGATAGCATTCATGCTGCCATGGGAAGTTGACCTCTCTCTAACAACAGGCATGGGCAGGAAGCTGACAGGATTAGTAGGGGCTGACACTGCTATCAGCCTTGGGGCACTTTGATCACAA
Mus_spretus         AAGATAGCATTCATGCTGCCATGGGAAGTTGACCTCTCTCTAACAACAGGCATGGGCAGGAAGCTGACAGGATTAGTAGGGGCTGACACTGCTATCAGCCTTGGGGCACTTTGATCACAA
Mus_caroli          AAGATAGCATTCATGCTGCCATGGGAAGTTGACCTCTCTCTAACAACAGGCATGGGCAGGAAGCTGACAGGATTAGTAGGGGCTGACACTGCTATCAGCCTTGGGGCACTTTGATCACAA

Mus_musculus        GGCTGGGAGAGAGGAATCTCACTGGACTTTTTTT---TTTTTTTTAACCTGAGCTAGGAAATCTTCCATCTCCTCTTTATTAACTTGGAATTGGGTCATCTGTTACTTCAGCTAACGGGA
Mus_spretus         GGCTGGGAGAGAGGAATCTCACTGGACTTTTTT------TTTTTTAACGTGAGCTAGGAAATCTTCCATCTCCTCTTTATTAACTTGGAATTGGGTCATATGTTACTTCAGCTAACGGGA
Mus_caroli          GGCTGGGAGAGAAGAATCTCACTGGACTTTTTTTTTTTTTTTTTTAACGTGAGCTAGGAAATCTTCCATCTCCTCTTTATTAACTTGGAATTGGGTCATCTGTTACTTCAGCTAACGGGA

Mus_musculus        TCCCTGGTGGAAACCTCGTTGGAAGAGCTGTTATGAGACCCACTAGGAAAGCATGTGACCGTCGCCTGCTAATTAAAAGGCTTGCCACTGAAATAATCAGCAGGCACATGCCCCTGGCAC
Mus_spretus         TCCCTGGTGGAAACCTCGTTGGAAGAGCTGTTATGAGATCCACTAGGAAAGCATGTGACCGTCGCCTGCTAATTAAAAGGCTTGCCATTGAAATAATCAGCAGGCACATGCCCCTGGCAC
Mus_caroli          TCCCTGGTGGAAACCTCGTTGGAAGAGCTGTTATGAGATCCGCTAGGAAAGCATGTGACCGTCGCCTGCTAATTAAAAGGCTTGCCACTGAAATAATCAGCAGGCACATGCCCCTGGCAC

Mus_musculus        ACGGATTCCAGGGAAATGCAAGCCTTTCATCTCTCTTCCATAGTATCTT-TTTTCAGTTGTAAATTCTTGGTTCAGTTTGCTTGAAATTCCCAAGCTACATAAGAAATAACTTAGTAGAA
Mus_spretus         ACGGATTCCAGGGAAATGCAAGCCTTTCATCTCTCTTCCATAGTATCTT-TTTTCAGTTGTAAATTCTTGGTTCAGTTTGCTTGAAATTCCCAAGCTACATAAGAAATAACTTAGTAGAA
Mus_caroli          ATGTATTCCAGGGAAATGCAAGCCTTTCATCTCTCTTCCATAGTATTTTTTTTTCAGCTGTAAATTCTTGGTTCAGTTTGCTTGAAATTCCCAAGCTACATAAGAAATAACTTAGTGGAA

Mus_musculus        AGAAAAGGAAGAAAACTGTGAGAAAACTACAAGTCTCTCTCTCTGGGCCTCTTCTTTTCTTTTTCCCCTCCCTCTTTCATTTCCTCTCACACACACTGGCCTCTAGCTCACTGGTCTCTT
Mus_spretus         AGAAAAGGAAGAAAACTGTGAGAAAACCACAAGTCTCTCTCTCCGGGCCTCTTCTTTTCTTTTTCCCCTCCCTCTTTCATTTCCTCTCACACACACTGGCCTCTAGCTCACTGGTCTCTT
Mus_caroli          AGAAAAGGAAGAAAACTGTGAGAAAACCACAAGTCTCTCTCTCCG-GCCTCTTCTTTTCTTTTTCCCCTCCCTCTTTCATTTCCTCTCACACACACTGGCCTCTAGCTCACTGGTCTCTT

Mus_musculus        ACCTCAGCCTCCTGGGTGCTGGGATTACAGGCCCAAGCCACTACGTCCAAGTTTGACTATCAAATTTCTTCCAGCATAATGTTGGCTCTTTCAAAGATTCTTTTCCCTCTCTCCCATTGT
Mus_spretus         ACCTCAGCCTCCTGGGTGCTGGGATTACAGGCCCGAGCCACTACGTCCAAGTTTGACTATCAAATTTCTTCCAGCATAATGTTGGCTCTTTCAAAGATTCTTTTCCCTCTATCCCATTGT
Mus_caroli          ACCTCAGCCTCCTGGGTGCTGGGATTACAGGCCTGAGCCACTACGTCCAAGTTTGACTATCAAATTTCTTCCAGCATAATGTTGGCTCTTTCAAAGATTCTTTTCCCTCTATCCCATTGT

Mus_musculus        GGATGACATCATGGAAAGAGTGTGTCGGAGCTAGCCAGACAGGAAGCCTGAGAGACAGGCCCAGGCATGTACCTTTATAAAAACACTCTCTCAAGAGCCCAACCAGGG
Mus_spretus         GGATGACATCATGGAAAGAGTGTGTCGGAGCTAGCCAGACAGGAAGCCTGAGAGACAGGCCCAGGCATGTACCTTTATAAAAACACTCTCTCAAGAGCCCAACCAGGG
Mus_caroli          GGATGACATCATGGAAAGAGTGTGTAGGAGCTAACCAGACAGGAAGCCTGAGAGACAGGCCCAGGCATGTACCTTTATAAAAACATTCTCTCAAGAGCCCAACCAGGG
```

**Supplemental Figure 6: -21kb *Igf2r* enhancer CLUSTWAL alignment**

Evolutionary conservation within -21kb *Igf2r* enhancer. Black boxed sequences indicate E/D boxes.

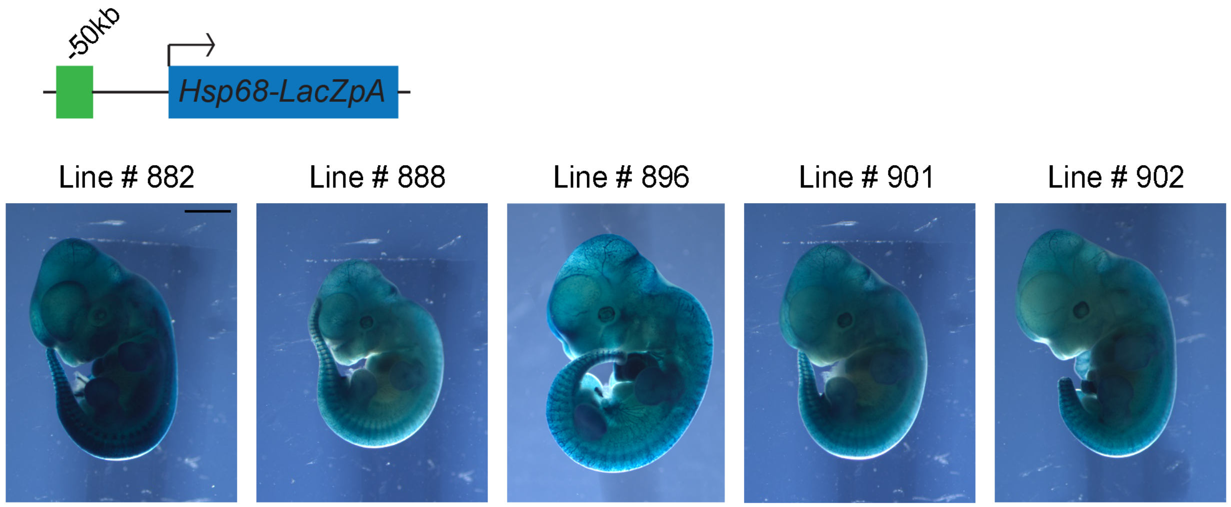


**Supplemental Figure 7: *LacZ* stable transgenic lines of -50kb *Klf2* enhancer**

Representative images of stable -50kb *Klf2* conserved non-coding element *lacZ* transgenic lines at E11.5. Scale bar 1mm.

**Supplemental FIgure 8**

```
Mouse AAGGGCCAGATGTGCTGAAACAGAACTTCAGTCCCAGCACCAGGGAGGCAGAGACACAGGCAAAGGCAGCTGAATCTTTATGAGTTCAAGGCCAGTCTGG
Rat   AAGGACCAGGTGTGCTGGCGTAGGCCTTCAGCGCCAGCACTAGGGAGGCAGAGACACAGGCAGAGGCAGGTGGATCTCTGTGAGTTCGAGGCCAGCCTGA
Human GTGGGCTGGTTGCGGTGGCTCACACCTGTAATCCCAGCACTCGGGAGGCTGAGACAGGA---GAATCACTTGAACCT--GGGAGGTGGAGATGAGGC---
Cow   GAG-------------------------------------------------------------------------------------------------

Mouse ACTACAAATTGAGAGCAAGGACAGCCAAGGCTACACAGAGAAAT-CCTATCTCAAAAAAAAAAAGAAAGAAAGAAAGAAAAGAAAAGAAAAAGAAAGAA
Rat   ACTACAAATCAAGATCAAGGACAATCAAGACTTCACAGAGAAAC-CCTGTCTCAAAAAAAACAAAGGGAAAAAAAAACATT-------------------
Human ----------AATATCGCCTCCAGCCTGGGTG--ACAGAGCAAG-AATCTATCAAA-----------------AAAAGAGAGACCCAAACCAACAAAAAA
Cow   ------------------------------------------------------------------------------GGAGATGAAAGCCAGCG-----

Mouse AAGGAAAAAACCCGAGTTTT-TAAAATAAAAGAAC-AAT-ATTCTGAGCCTGGTGATGTGGCGATGTGGTGT-GTGCC--CG-CGCTGGGGAGGCTGAGG
Rat   -----------------TTT-TAAAATAAAAGAAT-AAT-ATCCTGAGCCTCGTGATTTG--------GTAT-GTGGT--GG-AGCTGGGGACGCTGAGG
Human GAGAGA-----GAGAGAATCACAATGTAAGTGAAT-AGT-GTAATTAGCCAGGCATGGTGGCAGATGCCTGTGGTCCC--AGCTACTTGGGAGGCTGAGG
Cow   -------------GAGAATCAGA------GTGAAA-TGA-GTA---------------------------------------------------------

Mouse CAGGAGAC-TG-ACTTCAGGCC-GCTTGGGCCACA-GAACAAGGCCCTAGGTCAAAACCACAACCACAACAATAGCAAAGGAAT-TCTCCAAAAGTCT--
Rat   CAGGAGAC-TG-ACTTCAGGCTATCTCTGGCCACA-GAACAAGGCCCTAGGTCAAAACCACAACCACAACAATAGCAAAGGAAT-CCTCCAGAAGTCTGT
Human CAGAAGAA-TA-GCTTCACTCCAGCCTGGGCAACAAGAGCAAGACTCCATCTCAAAAAAAA----------AAAAAAAAGGAAT-CTTCCGGAAGCTGAG
Cow   -----------------------------------------------------------------------------------T-CTCCCAGGAGCTCAG

Mouse GTCTTGGTCAGTGAGGGCAGAGGGGAAAAATCAGATTTCCAGGGGGCGAGAGGGCTCTGTGGGTGCTTGATGCCAAGTCTTAGGACCAGGACCCACGTGG
Rat   GTCTTGGTCAATGAGGGCAGGGGGGAAAAA-CAGATTTGGAGGGAGCGAGAGGGCTCTGTGGGTGCTTGCTGCCAAGTCTTAGGACCAGGAC--------
Human ATCTTGGTCAGCAGGTGCAGAGG-AAAGAGCCAGGTTTCAGAGCCAC-----------------------------------------------------
Cow   GTCTTGGTCGGCAGATCTAGGAG-GAAGAATCAGGTTTCAGAGCCAT-----------------------------------------------------

Mouse TAGAAGAAGAGAGATGACTCCCCTGCGTTGCCCTGTGACTTCTCCATGTACACCGTGGCATAAGGACATGCACTCACACAAATTTTAAAAGTTAAAAAAA
Rat   TGGAAGAAGAGAATTGACTCCCCCACGCTGCCCCGTGACTTCTCCATGTACACTGTGACATGTGCACATGCACTCATACAAATTTTAAAAATTAAAAAAA
Human ----------------------------------------------------------------------------------------------------
Cow   ----------------------------------------------------------------------------------------------------

Mouse AAAAAAAACTTTAAAGAATCAGACTTTCAGACTTAGAGCTCCATCTTATAACCTCCATTTCCTCCTCTGGGAAATGGGGACAGTAGAAGCTTGACTCTGA
Rat   -AAAAGAACTTTAAAGAATCAGACTTTCTGACTTAAAGCCCCATCTTGTAACCTCCATTTTCTCCTCTGGGAAGTGGGAACAGTAGGAGCTTGACTCCGA
Human --------------AGAATCA-----------------GTTTCTCTTGGAATCTCAGGTTCCTTCTCTGTGAAATGGGGATAGTGACACTCTGACTCCAA
Cow   --------------GGAATCA-----------------CCCCCTCTTGGAATTTCAGCTTCCTTCTCTGTGAAATAGGGAGAGTGTCACT-TGACTCCAA

                                                                                        Ebox1
Mouse GCATTCCCACATG-TGTTTCTTTAGCCTGGTCACCCCTTCCAAACGATCTTCCTGGTGAACTTCCGT----GAAACTTCACCTGCAGTGACCCTCCTCCA
Rat   GCATTTGCACTTG-TGCTCCTTTAGCCTGGTCACCCCTTCCCAGCGATCTTCCTGGTGAACTTCTGTCTGGGAAACCTCACTTGCAATGTCCCTCCTCCA
Human GCATTCACACATGCTGTTCCCTCAGCGTGGTCATCCCTTCCGTGTCATTTGCCTGGAGAGCTCCTACATATTAAACCCCAACTGTGATGTCCCTCCTCCA
Cow   GCCTTCACATACACTGTTCCCTCAGCCTGGTCATCCCTTCCATGCCTTCTGCCTGATGAACTCCTACATATTAAACCCTAGCTATGACACCATTCCTCCA
```

```
                                            Ebox2
Mouse GACACTTTTGTGTTTCAGGGCCGGGGACATCTGTGTGCCCCTGGACTTGGAGCCCCTT---GA---------TGGACAC--AGAGGGTTGG-AGAGTGGT
Rat   GACACTTTTGTGCTCCAGGGCCAGGGACATCTGTATGCCCC-AGACTTGGAGCCCCTT---GA---------TGGACAC--AGAGGGTTGG-AGAGCATC
Human GGCAGCTCCAAACTGCAG-GCTGTGGCCATCTGTGTCCCCTGAGACTGGGGGCTCCTTGAGGACTGGGCTAGGGGACAC--AAGGGGATGG-CGAGTGTT
Cow   GGCAGTCCTGGACTGTAGGGCTG--------TGTGTGCCCCCAGACTGGAGGCTCCTT---GA---------TGCCCAGGCTGGGGGGTGG-GGAATGTT

Mouse AA-AGCAGAGACATCCAGGGCTCAGCAGTATGGCA-CGGGACTCCGAATACAGTGTTCTTTCCCAAAATCTGTGTGC--ACGAAA---GTGCCCAGCCAC
Rat   AA-AGCAGAGACACCCGGGTCTCAGCTCCACAGCA---GAACTCCGAATACAGTGTTCGTTCCCAAAATCTGTGCGC--GAGCCA---GTGCCTAGTCAC
Human AAAAATGGAGATGCCAGATTTTCAACTCTTGGGCAGCCAGACTGCAAGTCCAGTGCTCCTTCC-AAAACTTGTGTGT--ATGG-A---GAACCCA-----
Cow   AA-AATGAAGCC-----CAGTTTCAC-TTGGGGCAGTCGGGCTCCA---------CTCCATCTCAAAACTTGAGAGTGGACCTAA---CTACCCA-CCAC

Mouse GGTCAT----CTG--GGCCCTGCCCCC-TCCCC-ATCCTCACAGCCCCTGC-CAGGG--CCAGGACAGCCTGTCCCCTGAGT-GTCCCCCAGGTCATAGA
Rat   AGTCAT----CTG--GGCCCTGCCCCCATCCCC-ATCCTTACAGCCCCTGC-CAGGG--CCAGGACAGCCTGTCCC--GAGT-GTCCTCCAGGTCACAGA
Human --CCAC----CTG--TGCCCTGACCCCCTCCTGGTTCCCCACAGCCCCTGC-CCAGG--CCAGGACACCCTGTCCCCTGGGT-GTCCCTGAGGTCACAGA
Cow   GCTCAGAGGGCTGCTCGCCCTGACCCCCTCCTGGGTCCCTGCAGCCCCTTCCCTGGGACCTAGAACAGTCTGTCCCCTTGGT-GGCCCCAGGGTCACAGA

                                                                 Ebox3
Mouse TAAGGCTGCCAG-TGG-CTATCGGCAGGAAACATCTTAGCAAG-CAGGAAACCACGAAGTGAGCCAACTGCGCCCGCGCCGAGCTCAGCCTATCCTTGGC
Rat   TAAGGCTGCCAG-TGGGCTATCGGCAGGAAACATCTTAGCAAG-CAGGAAACCACGATGTGAGCCAATTGCGCCCGCGCAGGGCTCAGCCTATCCTTGGC
Human TAAGGCCGCTGGGCCGGCTATCGGCAGGAAACATCTTGGCAGG-ACAGGAAGCGCGGCATGGGTCAGCCGCGCCTGCACCCGGCTCAGCCTATCAGCGGC
Cow   TAAGGCCACTGGGCCAGCTATCAGCAGGAAACATCTTGGCAGGACAGGAAGCCACGGCATTGGCCTGCCGCACCCTCAGCCGCCTCAGCCTATCCACGGC

Mouse CTGATATTTTGGGAGCCGCTATTTAAGC-CCGACTCGACTCCTGCCATGAGCTGGACTGTTCTTGTGAGGTTGGAGGGAGACTCCATTTCCACATCCCTC
Rat   CTGATATTTTGGGAGCCACTATTTAAGC-CCGACTCGGCTCCTGCCCTGAGCTGGACCATTCTCCTGGAGTTGGAGGGAGGCTCTAGTTCCACATTCCTC
Human CAGATATTTTGGGAGCCACTATATAAACCCCAACTCGGCTCC------AAGC-----TGTTCCCCG--------GGGAAGCCTCAGGCTTCATGTTCTGC
Cow   CTGATATTTTGGGAGCCGCTATATAAACCCTGACTGGGCTGC------CAGT-----TGTTCCCGG--------GGGAAGCGCGAAGCTTCACGTTCTTC

Mouse AGAAG-CC--------TGCGGC--CAGAGATATGAGGATCGGA-----TCACAGA-GGCCC------TCTAGAGAGCC-----ACAG--CCTCTTCTCTA
Rat   ACAAG-CC--------TGAGGC--CAGAGATACGAGGAATGGA-----TCACAGA-GAC-C------TCTAGAGAGTC-----AGAG--TCTCCTGTCTA
Human GGCAGCCC--------TGAGGCTGCTGGAACACGG-GGCCCGGGAGGAGCTCAGT-AG-CC------CCTATTCAGTCCTGGTGCAGA-CACTCTCACTT
Cow   GGCAGCCCTGTGGCCCCTGGGC------------------AAG---ATGCTCAG-----TCCCCCGCCCCATTCAATCTCAGTGC-AC-C------ACTG

Mouse GCC-GCTGGGCAGCCC-GAGCATGTGGCGCCAGTTTT--CCCCTTC--ACAGCTTGGATTCTGTCCTAA---ATGAAAAACC----AAGTAGGAGGCCAG
Rat   GCTTGCTGGGCAGCCC-GAGCAGGTGGCTACAGTTTTTCCCCTTTC--ACAGCTTGGATTCTGCCCTAA---ATGAAAAACC----AAGCAGGGGGCCAG
Human ----GC---------T-GGGCAAGTGGCTCCAGTTTC--CCCCTCCCCACACCTTTGATCCTGTCCCAG---GGAA-AAACA----CACC-----TCCAG
Cow   GCTGGCTGAGCGACCCAGGGCAAACAGCTCTGGTTTT--TCCGTCCCCATACC-CCGGTCCTGTCCCAG---GA--AACACA----CTCACGGGTTCTAG

Mouse GT-CATGCCTGAAC-CAGCCTGTGAGA-GAGACCCATGTTCAA-GCAGTGCAC------A-TACATGCACACACAATATAAAGTAAATAAAAATTGGGG
Rat   AG-CATGTCTGAAG-CAGCCTGTGAGA-AAGACCCACGTTCAA-GCAGTGCAC------A-TACAGGCATGCACAATATAGAGTAAATAAAAATAGGGG
Human GT-CCTTTCTGAAC-CAGCCTGGGCAATAAGA------------GCAAGACTCCGTCTAAACACACACACACACACACAACA-------------------
Cow   AG----TCCTGAAC-TAGCCTC-------------ATA--------------------------------------------------------------
```

**Supplemental Figure 8: -50kb *Klf2* enhancer CLUSTWAL alignment**
Base pairs in light blue show regions of evolutionary conservation within -50kb *Klf2* enhancer. Black boxed sequences indicate E/D boxes. Ebox 1, 2, and 3 assayed by ChIP are highlighted in green and marked.

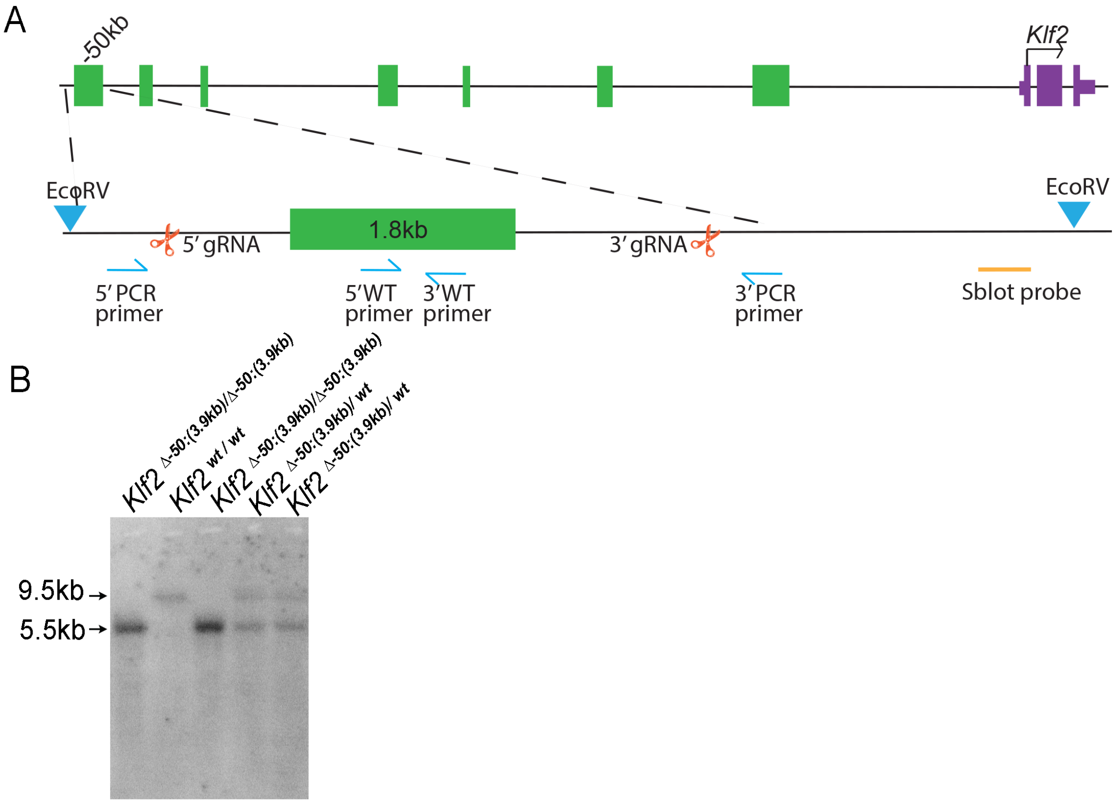


**Supplemental Figure 9: Design and validation of the *Klf2*$^{\Delta\text{-}50kb(3.9kb)}$ gene edited allele**

**A.** Deletion of -50kb *Klf2* enhancer by CRISPR/Cas9 with 5'- and 3'- guide RNA (gRNA, red scissors). Total size of deletion is 4300bp exceeds the size of the identified enhancer and was necessary to find efficacious gRNAs. PCR primers used for genotyping (in blue), wildtype product size 450bp, deletion product size 350bp. *EcorV* restriction digest (blue triangles) are used to generate RFLPs for Southern blots.

**B.** Southern blots for the wildtype (*Klf2*$^{wt/wt}$) RFLP migrate at 9.5kb and successful gene-edited enhancer deletion mutants (*Klf2*$^{\Delta\text{-}50kb(3.9kb)/\Delta\text{-}50kb(3.9kb)}$) RFLPs migrate at 5.5kb. Heterozygous alleles (*Klf2* $^{\Delta\text{-}50kb(3.9kb)/wt}$) contain both RFLPs.
